# Neuronal Tau Pathology Worsens Late Phase White Matter Degeneration After Traumatic Brain Injury in Transgenic Mice

**DOI:** 10.1101/2023.05.27.542567

**Authors:** Fengshan Yu, Diego Iacono, Daniel P. Perl, Chen Lai, Jessica Gill, Tuan Q. Le, Patricia Lee, Gauthaman Sukumar, Regina C. Armstrong

**Affiliations:** Anatomy, Physiology and Genetics; Neurology; Pathology; Department of Defense-Uniformed Services University Brain Tissue Repository; Center for Neuroscience and Regenerative Medicine, and; Henry M. Jackson Foundation for the Advancement of Military Medicine at the Uniformed Services University of the Health Sciences, Bethesda, MD; Johns Hopkins University, Baltimore, MD

**Keywords:** neurogeneration, traumatic brain injury, white matter, tau, demyelination, axon damage

## Abstract

Traumatic brain injury (TBI) causes diffuse axonal injury which can produce chronic white matter pathology and subsequent post-traumatic neurodegeneration with poor patient outcomes. Tau modulates axon cytoskeletal functions and undergoes phosphorylation and mis-localization in neurodegenerative disorders. The effects of tau pathology on neurodegeneration after TBI are unclear. We used mice with neuronal expression of human mutant tau to examine effects of pathological tau on white matter pathology after TBI. Adult male and female hTau.P301S (Tg2541) transgenic and wild type (Wt) mice received either moderate single TBI (s-TBI) or repetitive mild TBI (r-mTBI; once daily x 5), or matched sham procedures. Acutely, s-TBI produced more extensive axon damage in the corpus callosum (CC) as compared to r-mTBI. After s-TBI, significant CC thinning was present at 6 weeks and 4 months post-injury in Wt and transgenic mice, with homozygous tau expression producing additional pathology of late demyelination. In contrast, r-mTBI did not produce significant CC thinning except at the chronic time point of 4 months in homozygous mice, which exhibited significant CC atrophy (−29.7%) with increased CC microgliosis, but not astrogliosis. Serum biomarker quantification demonstrated neurofilament light detection of early axonal damage one day post-injury in Wt and homozygous mice. At 4 months, high tau and neurofilament in homozygous mice implicated tau in chronic axon pathology. Conclusions: Neuronal tau pathology differentially exacerbated CC pathology based on injury severity and chronicity. Ongoing CC atrophy from s-TBI became accompanied by late demyelination. Pathological tau significantly worsened CC atrophy during the chronic phase after r-mTBI.

## INTRODUCTION

Traumatic brain injury (TBI) can lead to post-traumatic neurodegeneration of gray and white matter brain regions that is associated with poor patient outcomes [9] [34]. More specifically, diffuse axonal injury in the corpus callosum is associated with post-traumatic neurodegeneration and unfavorable functional outcomes [34] [81]. Post-traumatic neurodegeneration can result from progressive secondary pathology following a single moderate-severe TBI or from milder repetitive concussive and subconcussive injuries. Secondary processes that contribute to post-traumatic neurodegeneration are important to identify and warrant high priority as therapeutic targets to mitigate the detrimental effects of TBI on long term brain health [12].

The purpose of this study was to test neuronal tau pathology as a contributing factor in the post-traumatic degeneration of white matter tracts that exhibit traumatic axonal injury, a hallmark pathology of TBI. Degeneration of cerebral white matter, including the atrophy of the corpus callosum, has long been recognized in postmortem cases after a single moderate-severe TBI [74] [44]. Recent magnetic resonance imaging (MRI) studies reveal that focal damage to densely connected white matter regions of brain networks results in worse cognitive impairment than similar damage to gray matter hubs [51] [62]. Furthermore, MRI parameters of white matter diffuse axonal injury predict subsequent white matter atrophy as well as whole brain atrophy over time after moderate-severe TBI [33]. Post-traumatic white matter atrophy involves deep white matter, e.g. corpus callosum and corticospinal tracts, which are associated with axonal damage [32]. A plasma biomarker of axonal damage, neurofilament light protein, correlates with MRI measures of diffuse axonal injury and predicts both chronic phase white matter neurodegeneration and functional outcomes after TBI [34] [59]. Brain atrophy, including reduced corpus callosum volume, and higher plasma neurofilament light levels have also been found after repetitive TBI in boxers [7]. Similarly, repetitive sports related head injuries diagnosed by neuropathology as chronic traumatic encephalopathy (CTE) exhibit post-traumatic neurodegeneration with cerebral atrophy that includes white matter changes of corpus callosum thinning, myelin rarefaction, axon loss, and altered glial phenotypes [3] [15] [19].

Neuronal phosphorylated tau pathology has been implicated in secondary injuries after TBI due to association with neurodegenerative diseases and being observed in postmortem cases of CTE as well as after single events of more severe TBI [19] [38] [69] [82] [84]. In the healthy adult brain, tau is localized in axons and serves as a microtubule associated protein that modulates axonal transport and stability of the axon cytoskeleton [70] [80] [90]. Tau localization in axons and binding to microtubules is regulated by phosphorylation [70] [90]. A range of genetic and acquired conditions can lead to tau hyperphosphorylation, mis-localization, aggregation and/or impaired clearance [11] [38] [40] [41]. Such findings indicate potential interplay between TBI sequelae and tau pathology in the progression of white matter degeneration that remains poorly understood.

In this study, we tested the effect of increasing levels of neuronal tau pathology on the progression of white matter degeneration after a moderate single TBI (s-TBI) or after repetitive mild TBI (r-mTBI) in transgenic mice expressing human tau with a pathogenic mutation. Varied pathology, particularly the extent of corpus callosum axon damage in these two different injury models, was tested relative tau transgene dosage to inform interpretation of heterogeneous patient populations. As extensively characterized in adult wild type mice, the s-TBI model produces traumatic axonal injury that progresses to corpus callosum atrophy, with features that model clinical findings of neuropathology and MRI of patients with TBI [10] [33] [53] [54] [61] [71]. Similarly, the r-mTBI model has also been previously characterized in wild type mice using neuropathology and MRI [88]. The r-mTBI model uses a milder impact separated by 24 hours across five consecutive days, which produces traumatic axonal injury at acute and subacute time points, but does not degenerate so far as to produce overt corpus callosum atrophy [88] [60]. Progressive tau pathology pertinent to human histopathology has been demonstrated in adult wild type mice in recent studies that also used a five-day repetitive mild head injury, along with interpretations from a systematic review of prior literature [45].

Prior studies in wild type and human tau transgenic mice have advanced our knowledge of the molecular mechanisms of tau pathology and informed our use of human tau transgenic mice for analysis of post-traumatic neurodegeneration [5] [31] [45] [82]. Supplemental Table SI-1 lists examples of specific studies that are particularly relevant to the purpose of the current study.

Although the hTau mouse line enabled expression of all six isoforms of human tau driven from the human tau promoter [4], only hemizygous hTau mice are viable and the level of human tau expressed did not alter the neuropathological sequelae of TBI [27] [56]. We use hTau.P301S (Tg2541) mice that offer multiple advantages for the current studies of phosphorylated tau pathology as contributing to post-traumatic neurodegeneration. Tg2541 mice express the P301S mutation of the human tau gene driven in neurons from the Thy1.2 promoter [2]. Using Thy1-YFP-16 mice, we previously demonstrated that neurons expressing a fluorescent reporter protein from the Thy1.2 promoter exhibit axon damage in the corpus callosum and cerebral cortex in both s-TBI and r-mTBI models, with the most prominent axon damage in the corpus callosum after the moderate s-TBI [53] [88]. The Tg2541 mice have more reproducible tau expression than hTau.P301S (PS19) mice, which have extensive variability in both the rate of tau spreading and phosphorylated tau pathology [83]. Interestingly, studies in Tg2541 mice revealed molecular interactions that regulate phosphorylated tau pathology in the adult forebrain [43], which may indicate the potential for modulation by TBI sequelae. Furthermore, recent phosphoproteomics analysis in Tg2541 mice identified protein co-expression modules that correspond with cellular responses to phosphorylated tau pathology [79]. These findings in Thy1-YFP-16 mice and Tg2541 mice support our current study design and inform future studies of the cellular and molecular basis of tau effects in TBI models.

In the current study, we first confirmed in the Tg2541 line that the moderate s-TBI produces more extensive axon damage based on neuropathology. We also validated expression of reproducible differences in human tau protein in brain lysates from regions under the impact site, i.e. the rostral corpus callosum and cerebral cortex, with significant gene dosage effects across wildtype, hemizygous (Hemi), and homozygous (Hom) mice generated from the Tg2541 colony. We then conducted clinically relevant neuropathologic and blood biomarker assessments of the Tg2541 genotype effect at acute (1 day), subacute (6 weeks) and chronic (4 months) phases after s-TBI and r-mTBI. This translational approach demonstrated that phosphorylated tau pathology, axon damage, neuroinflammation, demyelination, and corpus callosum atrophy progressed with distinctly different timing and extent based upon the injury parameters in combination with the human mutant tau gene dosage.

## METHODS

### hTau.P301S (Tg2541) Mice

All animals were treated in accordance with guidelines of the National Institutes of Health Guide for the Care and Use of Laboratory Animals. The study protocol was approved by the Institutional Animal Care and Use Committee of the Uniformed Services University of the Health Sciences. Mutant human tau transgenic mice (B6-Tg(Thy1-MAPT*P301S)2541), also known as hTau.P301S (Tg2541) or Tg2541 mice, were generated by Dr. Goedert [2] with breeders backcrossed with C57BL/6J mice, and generously provided by Dr. Tony Wyss-Coray (Stanford University, Palo Alto, CA). This hTau.P301S Tg2541 line expresses the Thy1.2 promoter driving neuronal transcription of the P301ShTau43 construct to express the 383 aa isoform of human 4R Tau with a P301S mutation (Supplemental Information Table SI-1). Hemizygous mice were bred at the Uniformed Services University of the Health Sciences to generate experimental mice. Mice were housed in standard cages in a 12-h/12-h light-dark cycle with food and water ad libitum.

DNA from mouse ear biopsies was used for genotyping with quantitative PCR using primer pairs for both the inserted vector sequence (5’-AAAGGAACTCAACTCCACCTCAC-3’ and 5’-ACCTTACTGAGCTAGCAGGTCTTT-3’) and for the mutant human MAPT 5’-(GATTGGGTCCCTGGACAATA-3’ and 5’-GTGGTCTGTCTTGGCTTTGG-3’). Additionally, early in establishing the colony, some mice were genotyped for HuMAPT mut detection using DNA from tail biopsies with analysis by Transnetyx (Memphis, TN) using primers pair of 5’-CAGGAGTTCGAAGTGATGGAAGA-3’ and 5’-AGCCCCCCTGATCTTTCCT-3’ with reporter CCCGTACGTCCCAGCGTG.

### Traumatic Brain Injury Procedures

Two different models of TBI were used to reflect different human injuries associated with tau pathology and post-traumatic neurodegeneration. Both male and female mice were used for sham or TBI procedures at 8 weeks of age. An Impact One Stereotaxic Impactor (Leica Biosystems, Deer Park, IL) was used to produce concussive models of repetitive mild TBI (r-mTBI) [88] or single moderate TBI (s-TBI) [53] [54] [75], as previously characterized to produce pathology in the rostral corpus callosum and medial cerebral cortex. Mice were anesthetized with 2.0% isoflurane in O_2_ and positioned in a stereotaxic frame with the head fixed with ear bars capped by rubber stoppers for s-TBI and r-mTBI procedures. The impacts were made at stereotaxic coordinates for bregma (0 ML, 0 AP, 0 DV) using a 3-mm diameter tip with a flat surface and rounded beveling of the edge (Cat# 2530-3S, Neuroscience Tools, Aurora, IL). For s-TBI, after depilation of the scalp hair with Nair, a midline incision was made to expose skull and impact was made onto the intact skull at bregma (velocity at 4.0 m/sec; depth of 1.5 mm; dwell time of 100ms). For r-mTBI, after shaving and depilation of the scalp hair, impacts (velocity at 4.0 m/sec; depth of 1.0 mm; dwell time of 200 ms) were made onto the scalp approximately over bregma. The r-mTBI consisted of 5 impacts with one per day at 24-hour intervals on 5 consecutive days. Sham mice underwent identical procedures to their respective TBI mice without receiving impacts. To mitigate potential pain experienced from the injury procedures, mice were given acetaminophen (1 mg/ml) in the drinking water starting the day before a TBI or sham procedure and returned to standard drinking water one day after the final procedure. Body temperature was maintained with a warming pad. After each procedure, the duration of apnea and the righting reflex were recorded. The righting reflex was measured immediately after removal from anesthesia and placement of the mouse in the supine position to righting to the prone position, which is a surrogate measure of transient alteration of consciousness after TBI.

### Immunohistochemistry and Histological Staining

Mice were perfused with 0.1 M phosphate buffer followed by 4% paraformaldehyde at 1 day, 6 weeks, or 4 months after TBI and sham procedures. Brains and spinal cords were dissected and further post-fixed by immersion in the same fixative overnight. Tissues were embedded in paraffin and then cut as coronal sections (5 µm). Immunohistochemistry was used to detect tau and cell type specific markers with antibodies listed in Table 1. Briefly, HT7 detected human Tau isoforms while AT8 immunolabeled tau phosphorylated at S202/T205. Microglia/macrophages were identified based on ionized calcium-binding adaptor molecule 1 (IBA1) immunoreactivity. Astrocytes were immunolabeled for glial fibrillary acidic protein (GFAP). Axon damage was evaluated based on beta-amyloid precursor protein (β-APP) or phosphorylated neurofilament H (SMI-34) antibodies. Myelination was immunolabeled for myelin basic protein (MBP). Immunohistochemistry was performed analogous to the human TBI neuropathology processes, using Leica Bond III automated staining system with diaminobenzidine chromogen detection and the same antibodies as for clinical diagnostics (DS9800, Leica Biosystems, Buffalo Grove, IL). Hematoxylin and eosin stain was used for general histopathology. Sections were scanned to create digital images with either NanoZoomer (Hamamatsu Photonics, Japan) or Aperio (Aperio AT2 - High Volume, Digital whole slide scanning scanner, Leica Biosystems, Deer Park, IL) digital slide scanning systems and images were captured and exported with NDP viewer (Hamamatsu Corporation) or Aperio ImageScope (Leica Biosystems), respectively.

**Table 1.** Antibody Information

| Application | Primary Antibodies | Secondary antibodies |
| --- | --- | --- |
| IHC | HT7 (mouse monoclonal) 1:150<br>Thermal Fisher Cat # MN1000<br>RRID: (AB_231465) | Secondary antibodies inside the<br>diaminobenzidine chromogen<br>detection system (Leica<br>Biosystems, DS9800) |
|  | AT8 (mouse monoclonal) 1:200<br>Thermal Fisher Cat # MN1020<br>RRID: (AB_223647) |  |
|  | Iba1(rabbit monoclonal) 1:200<br>FUJIFILM Wako Pure Chemical<br>Corporation Cat # 019-19741<br>RRID: (AB_839504) |  |
|  | GFAP (rabbit polyclonal) 1:100<br>DAKO Cat # GA524<br>RRID: (AB_2811722) |  |
| | β-APP (mouse monoclonal) 1:100<br>Millipore Sigma Cat # MAB348<br>RRID: (AB_94882) | |
|  | SMI34 (mouse monoclonal) 1:100<br>Biolegend Cat # 835503<br>RRID: (AB_2572004) |  |
|  | MBP (rat monoclonal) 1:100<br>Abcam Cat # AB7349<br>RRID: (AB_305869) |  |
| Wes | HT7 (mouse monoclonal) 1:100<br>Thermal Fisher Cat # MN1000<br>RRID: (AB_231465) | Goat Anti-Mouse Secondary HRP<br>Protein Simple Cat # 042-205<br>RRID: (AB_2860576) |
|  | AT180 (mouse monoclonal) 1:50<br>Thermal Fisher Cat # MN1040<br>RRID: (AB_223649) | Goat Anti-Mouse Secondary HRP<br>Protein Simple Cat # 042-205<br>RRID: (AB_2860576) |
|  | P-Tau (S404) (rabbit polyclonal) 1:1000<br>Thermal Fisher Cat # 44-758G<br>RRID: (AB_2533746) | Goat Anti-Rabbit Secondary HRP<br>Protein Simple Cat # 042-206<br>RRID: (AB_2860577) |
|  | P-Tau (T205) (rabbit polyclonal) 1:200<br>Thermal Fisher Cat # 44-738G<br>RRID: (AB_2533738) | Goat Anti-Rabbit Secondary HRP<br>Protein Simple Cat # 042-206<br>RRID: (AB_2860577) |
| | β-actin (mouse monoclonal) 1:50<br>Sigma Cat # A1978-200ul<br>RRID: (AB_476692) | Goat Anti-Mouse Secondary HRP<br>Protein Simple Cat # 042-205<br>RRID: (AB_2860576) |

Quantification of immunohistochemistry and histological staining was conducted with coronal sections from within approximately +0.5 to -0.5 mm of bregma (i.e. under impact site). Reactive astrogliosis and microgliosis were estimated from the area of immunoreactivity for GFAP or IBA1 using ImageJ (NIH, Bethesda, MD). The corpus callosum region-of-interest included both hemispheres extending laterally over the lateral ventricle and under the peak of the cingulum. The medial cerebral cortex was analyzed for the cingulate and motor cortex above the corpus callosum region [88]. Damaged axons in the corpus callosum or medial cerebral cortex were counted manually using NIH ImageJ (NIH) with β-APP or SMI-34 images, respectively based on the presence of large terminal end bulbs or as multiple swellings along a longitudinal axonal profile. Any β-APP or SMI-34 positive end bulb that was adjacent to or continuous with a nucleus was excluded. For quantification of axon damage, the hematoxylin nuclear counterstain was used to exclude immunolabeling associated with neuronal or glial cell bodies [28] [35]. Corpus callosum width was analyzed with hematoxylin and eosin staining or with immunolabeling for MBP. The width was calculated for each mouse from the average of 7 measurements (midline, and bilaterally approximately 200 µm from midline, under the peak of cingulum, and at the lateral border of ventricle [53] [54], using images digitized at 20 X on NDP viewer (Hamamatsu Corporation) or Aperio ImageScope (Leica Biosystems). HT7 immunolabeling was used to identify the presence or absence of human tau expression. Analysis of AT8 scored immunoreactivity, based on intensity and area, as negative or present (scale of + to ++++) to localize phosphorylated mouse or human tau in tissue sections as a complement to Wes quantification of tau protein in brain lysates (see below).

### Soluble Tau Protein and Phospho-tau Epitopes in Brain Lysates

Soluble tau in brain tissue was quantified with Wes Protein Simple Westerns™ (Protein Simple, San Jose, CA). Prior to tissue collection, mice were perfused with cold phosphate buffered saline (PBS) to flush out blood. The brain was cut rapidly with McIlwain Tissue Chopper (Stoelting Co., Wood Dale, IL) into 1 mm thick coronal slices and placed in cold PBS. The slice containing the anterior commissure was identified to estimate the slice aligned in the coronal plane with bregma. The upper forebrain was quickly dissected by a straight razor cut in the horizontal plane just below the lowest midline point of corpus callosum. This upper forebrain region under the impact site included the corpus callosum and cerebral cortex at coronal levels from two slices comprising approximately +1.0 mm to -1.0 mm relative to bregma. Tissue was stored in -80°C freezer. The soluble protein fraction was isolated with minor modifications of published protocols [20] [85]. Brain tissue samples were immersed in 3 X T-per buffers (#78510; Thermo Fisher, Waltham, MA) with protease and phosphatase inhibitors (#1861281; Thermo Fisher). After homogenization and sonication, the tissue suspension was separated at 80000g, 4°C for 30 minutes in an Optima^TM^ MAX Ultracentrifuge (Beckman, Germany). Supernatant was collected and stored at -80°C. Simple Westerns™ (Wes Protein Simple) were run according to the manufacturer’s protocol. Primary antibodies used and their conditions are listed in Table1. Data was analyzed with Compass software (Protein Simple). A peak on the electropherogram is accepted when the molecular weight is at the right position, and signal to noise ratio is higher than 10. All the peaks wider than 20 were excluded.

### Serum Biomarkers

Mouse serum was analyzed for biomarkers using the ultra-sensitive Single molecule array (Simoa ®) for the Neurology 4-Plex A multiplex assay (N4PA; Quanterix, Lexington, MA), as has been successfully used in clinical studies [29] [47] [49]. Cardiac blood was collected as a terminal procedure from the mice utilized in other experiments of this study. Blood was drawn from the right atrium of deeply anesthetized mice immediately before intraventricular perfusion. After remaining undisturbed at room temperature for at least 30 minutes, samples were centrifuged at 3000 g for 10 minutes. Serum were carefully collected and stored at -80°C until analyzed. Serum levels of tau, GFAP, neurofilament light chain (Nf-L) and Ubiquitin C-terminal Hydrolase-1 (UCHL-1) levels were simultaneously measured in each sample and samples were run in duplicate on a Simoa ® HD-1 Analyzer (Quanterix, Lexington, MA). The results with coefficient of variation values below 10% were accepted for final analysis. This assay is designed for detection of human proteins.

Performance with mouse proteins in serum was tested by the manufacturer (Quanterix application note MKT-0038 01). Immunodepletion tests demonstrated good specificity for mouse Nf-L and that the assay does not specifically detect mouse GFAP or UCHL-1 proteins, which is in agreement without our results (data not shown).

### Progression of Neurologic Impairment

In this line of hTau.P301S Tg2541 mice, phosphorylated tau pathology initiates in the spinal cord and mice develop corresponding paresis and paralysis with age [2]. Therefore, the hang time test was used to quantify early spinal cord dysfunction [26] [63]. Briefly, the mouse was placed onto a wire cage top held over the padded surface. The cage top was then inverted so that the mouse gripped the wire bars to support its own weight. The time hanging onto the cage top was recorded for up to 60 seconds. Mice with spinal cord pathology, including limb paralysis or paresis, do not have the grip strength to support their weight for 60 seconds and so drop to the padding at different time intervals depending upon the severity of the neuromuscular effect. Healthy adult C57BL/6 mice can hold on for the maximum trial time of 60 seconds without any difficulty, while mice that do not hang onto the cage bars for at least 30 second are considered impaired [26]. Hang time scores were calculated as the average of three hang time trials conducted in series with 15 second intervals to rest. After the hang time testing, mice were also observed for signs of limb paresis and paralysis with a neurological deficit score of 1 assigned for each affected limb up to a total score of 5 (limbs and/or tail).

### Mouse Numbers and Data Analysis

The experiments involved a total of 636 mice, which required different tissue collection so that 468 mice were perfused for immunohistochemistry and168 mice were prepared for protein quantification of brain lysates. Among these mice, 69 non-injured (naïve) mice evaluated with the hang time test to select post-injury time points relative to the progression of neurological deficits with the tau phenotype, 150 had behavioral testing based on sociability impairment, and cardiac blood was collected from 491 mice as part of the terminal procedure for the tissue analysis.

Experiments included approximately equal numbers of male and female mice based on the combination across cohorts of experimental mice available in litters from hemizygous breeders. Mice were allocated to sham or injured to distribute available mice to both conditions and balance sexes, to the extent possible within each litter. Most litters did not have sufficient mice for further steps to randomize within a sex and condition. Mice were allocated to injury or sham conditions prior to 8 weeks of age, which precedes behavioral signs associated with the tau phenotype. Mouse weights did not vary due to genotype at 8 weeks of age (data not shown). Experiments for the single impact TBI and corresponding sham condition have a visible scalp incision while the repetitive TBI and corresponding sham condition do not have a scalp incision; therefore, experiments in each model were conducted separately and were not randomized within cohorts.

Mice were excluded if the injury procedure inadvertently produced a depressive skull fracture, or if the mice had poor health to require euthanasia or otherwise died prior to the experimental endpoint. Humane euthanasia was required for mice that developed severe paraparesis. Power analysis for tissue section analysis was estimated from prior studies. However, the tissue analysis as defined by specific region of interest was important to maintain, so mice had to be excluded if sufficient tissue was no longer available. The Cohen’s d effect size is provided for the major findings of the neuropathology using the myelin basic protein immunohistochemistry results that had a low sample number. Analysis was conducted for each individual mouse for each condition so that values for each mouse are shown by symbols on the graphs for each experiment. All assessments were performed with blinding as to the mouse injury and tau genotype.

All data analysis was performed with GraphPad Prism 8 software (GraphPad Software, Inc., La Jolla, CA). Two-way analysis of variance (ANOVA) with post-hoc Holm-Sidak’s multiple comparisons test was used for statistical comparison of tau genotype and injury effects for the data from neuropathology of tissue sections, Wes Protein Simple and serum biomarkers. Neurological impairment based on the hang time test was analyzed as a Kaplan-Meier plot with the Mantel-Cox log rank test for statistical comparison. The full set of statistical comparison data is provided as tables in the Supplemental Information that are referred to in each figure legend. Statistically significant post-hoc comparisons are also shown on the graphs for each experiment. Statistical significance was defined as p < 0.05. This study did not generate big data resources but data from specific experiments will be made available upon request and statistical data from the current experiments is provided as Supplemental Information.

## RESULTS

### Acute injury severity is greater in s-TBI than r-mTBI at 8 weeks of age and is not increased by tau genotype

The study design took advantage of the distinct progression of neurologic impairment with hTau.P301S gene dosage in the Tg2541 mice (Figure 1). Homozygous mice are expected to develop overt neurologic symptoms of muscle weakness, tremor, and severe paraparesis at 5-6 months of age with a similar phenotype in hemizygous mice by 12-14 months [2]. We quantified the onset of neurologic symptoms with aging in non-injured (naïve) mice using a hang time test of grasping and strength (Figure 1A). In homozygous mice, hang time deficits developed by 4-6 months of age and progressed to paraparesis by 7-8 months of age. Based on this neurologic progression, the study was designed for mice to receive head injuries at 8 weeks of age with survival for 1 day (acute), 6 weeks (subacute), and 4 months (chronic) after injury or sham procedures (Figure 1A, B). This time course targeted analysis at the acute and subacute phases prior to the onset of symptoms and at the chronic phase after symptom onset indicative of progressive pathology in homozygous mice for comparison with hemizygous and wild type littermates.

**Figure 1.**
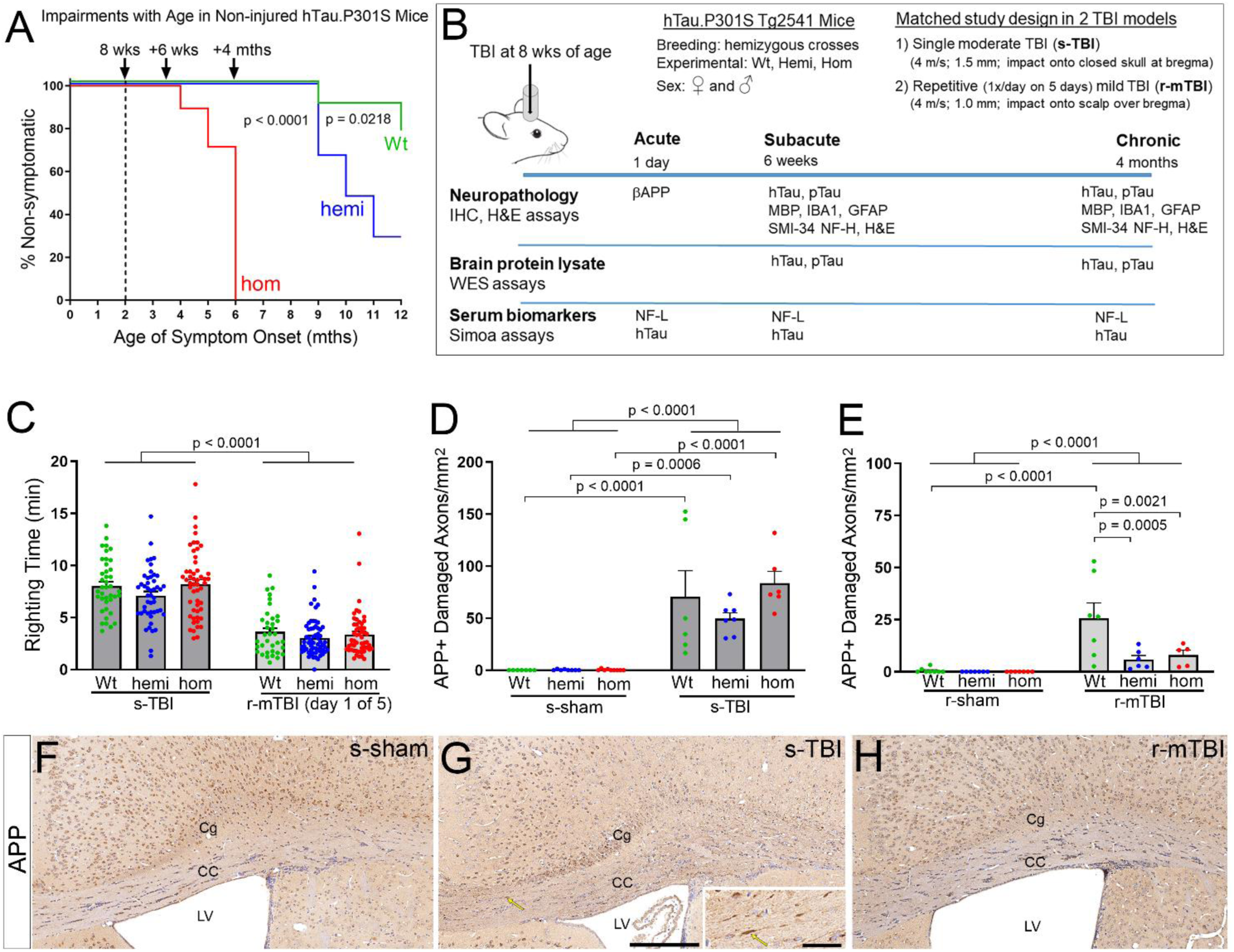
Acute injury severity is greater in s-TBI than r-mTBI and is not increased by hTau.P301S genotype at 8 weeks of age, prior to tau induced neurologic impairment. A. Hang time testing data. The ability of non-injured (naïve) mice to support their weight while hanging from wire cage top bars reveals progressive neurologic impairment with increasing age that is accelerated in homozygotes. This time course in naïve mice was used to select the time points for analysis after TBI. B. The study design matches mice across genotype, sex, and TBI models for differential comparisons at phases prior to and after symptom onset. Three distinct approaches examine tau expression for localization, quantification, and translational biomarkers. C. Immediately after TBI or sham procedures, a surrogate measure of loss of consciousness is an indicator of more severe injury after s-TBI compared to r-mTBI across genotypes. D. At 24 hrs after a single moderate TBI (s-TBI), acute axon damage in the corpus callosum is increased across genotypes. E. At 24 hrs after the last impact in the repetitive mild TBI (r-mTBI), the acute axon damage is reduced relative to s-TBI (Figure SI-2) and highest in wild type mice. C-E. Dots represent individual mice with wild type (Wt) in green, hemizygous (hemi) in blue, and homozygous (hom) in red. Non-significant comparisons are not shown. Full statistical analysis provided in Table SI-2, 3, 4. F-H. Representative images of β-APP immunolabeling in the corpus callosum (CC) area under the impact site in wild type mice from the Tg2541 line. Damaged CC axons were identified as β-APP immunolabeled swellings in longitudinal axon profiles (G, yellow arrow enlarged in inset). After s-TBI, damaged axons are also evident in the cingulum (Cg) as β-APP accumulated in transverse axon profiles. Hematoxylin staining of nuclei (blue) was used to exclude β-APP associated with neuron and glial cell bodies. LV = lateral ventricle. Scale bars: F-H shown in G = 200 µm, inset = 50 µm.

This time point of 8 weeks of age at the time of injury is matched to our prior studies characterizing the distinct s-TBI and r-mTBI injury models using neuropathology, electron microscopy, MRI, and behavior analyses [10] [53] [54] [61] [88]. The acute phase post-surgical and neuropathology data from these prior studies are similar to the results in the Tg2541 mice across wild type, hemizygous, and homozygous genotypes. Parameters of acute injury severity were increased in the moderate s-TBI model as compared to the r-mTBI model (Figure 1C-E).

Immediately after the injury or sham procedures, the s-TBI produced a longer delay than the r-mTBI in righting reflex, which is a surrogate measure for loss of consciousness (Figure 1C). The delay in righting time was significantly increased in injured mice relative to the sham condition in both the s-TBI and r-mTBI models, and was not altered across tau genotypes (Figure 1C and SI-1). The delay for s-TBI is compared with the first of the five days of r-mTBI impacts since this first day consistently produced the longest righting reflex time interval (Figure SI-1). Acute axon damage was quantified at 24 hrs after the s-TBI impact or the last of the five r-mTBI impacts using immunolabeling for β-amyloid precursor protein (APP), which detects early stage axon damage associated with impaired axonal transport, axonal fragmentation, and formation of terminal end bulbs (Figure 1D-H). As compared to the sham condition, significant CC axon damage resulted from s-TBI (Figure 1D, F, G) and r-mTBI (Figure 1E, H). Acute axonal injury was significantly more extensive after s-TBI as compared to the r-mTBI in mice of all three genotypes (Figures 1D-E and SI-2). Furthermore, the tau genotype did not increase acute vulnerability to axon damage with s-TBI or r-mTBI at 8 weeks of age (Figure 1D-E). The lack of a detrimental effect of tau genotype after acute injury is an important finding for attributing tau effects at subacute and chronic time points in our subsequent experiments.

### Cortical axon damage is increased at subacute and chronic stages in tau homozygous mice

A specific increase in axon damage due to tau genotype was evident as homozygous mice aged to subacute and chronic stages (Figure 2). While β-APP was effective for detecting acute stage axon damage (Figure 1), for longer survival time points we chose SMI-34 immunolabeling (Figure 2), which is advantageous for detecting axon loss and damage during subacute and chronic stages [30] [52] [76]. In healthy brain, neurofilaments are predominantly non-phosphorylated in neuronal cell bodies and dendrites, in contrast to the phosphorylated neurofilaments of axons [73]. SMI-34 immunolabeling of phosphorylated neurofilament heavy chain (NF200) detected beading and end bulb formation in damaged cortical axons, along with evidence of accumulated phosphorylated neurofilaments in cortical neuron cell bodies (Figure 2E-G). Axons in the corpus callosum were well labeled by SMI-34 in homozygous mice at 4 months after the sham surgical procedure (Figure 2H). However, at 4 months post-injury both s-TBI and r-mTBI homozygous mice (Figure 2I, J) had variable, mottled SMI-34 immunolabeling in the corpus callosum that may indicate axon loss or dephosphorylation, which is associated with neurofilament compaction in damaged axons [76] [78].

**Figure 2.**
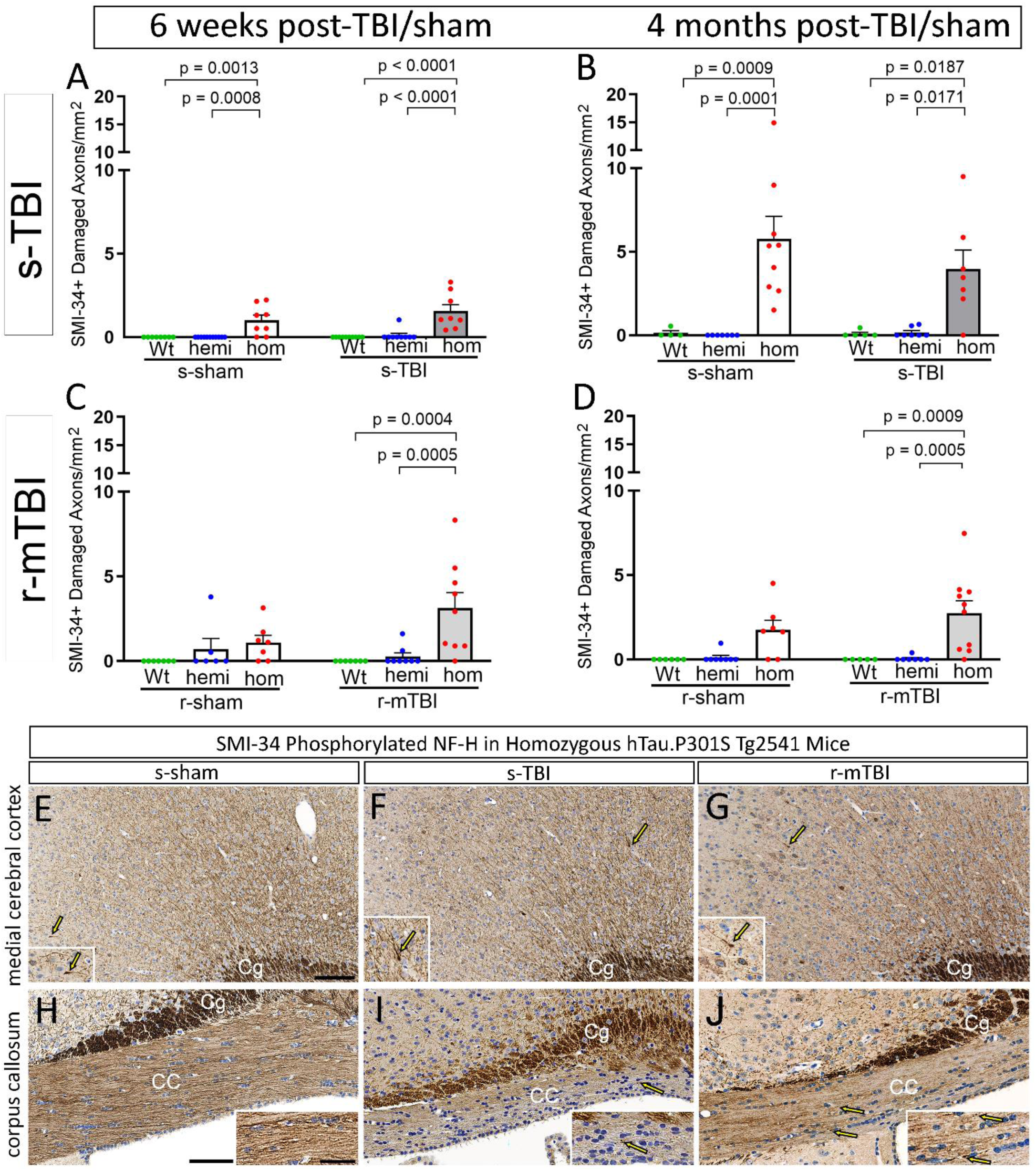
Homozygous hTau.P301S genotype increases cortical axon damage at subacute and chronic stages. A-D. Damaged axons with swellings or endbulbs identified with SMI-34 phosphorylated neurofilament immunolabeling in the medial cerebral cortex under the impact site at 6 weeks (subacute) or 4 months (chronic) stages after TBI or sham procedures. Dots represent individual mice with wild type (Wt) in green, hemizygous (hemi) in blue, and homozygous (hom) in red. Non-significant comparisons are not shown. Full statistical analysis in Table SI-3, 4. A-B. Axon damage is significantly increased in only homozygous mice after s-TBI or the s-sham procedure, which involves anesthesia and scalp incision. C-D. Axon damage is increased only in homozygous mice after r-mTBI, but is not increased after the r-sham procedure that involves only anesthesia. E-J. Representative images of SMI-34 immunolabeling in regions under the impact site in coronal brain sections from homozygous mice at 4 months after TBI or sham procedures. Yellow arrows indicate examples of damaged axons in the medial cortex and corpus callosum (CC) that are enlarged in each inset. Axons cut transversely in the cingulum (Cg) are strongly immunolabeled. Nuclei stained blue with hematoxylin. Scale bars: E-G shown in E = 100 µm, H-J shown in H = 100 µm, insets shown in H = 50 µm.

### Chronic stage phosphorylated tau neuropathology varies with head injury conditions and hTau.P301S genotype

We next evaluated tau pathology in coronal brain sections at the chronic stage of 4 months after s-TBI or r-mTBI, and corresponding sham conditions, and found injury induced phosphorylated tau in wild type mice that was amplified in homozygous mice. The initial characterization of the hTau.P301S Tg2541 line tested a battery of antibodies and reported that AT8 detected tau phosphorylation in the largest number of cells [2]. Therefore, we evaluated AT8 immunolabeling in medial cerebral cortex and corpus callosum regions under the impact site (Figure 3 and Table 2).

**Figure 3.**
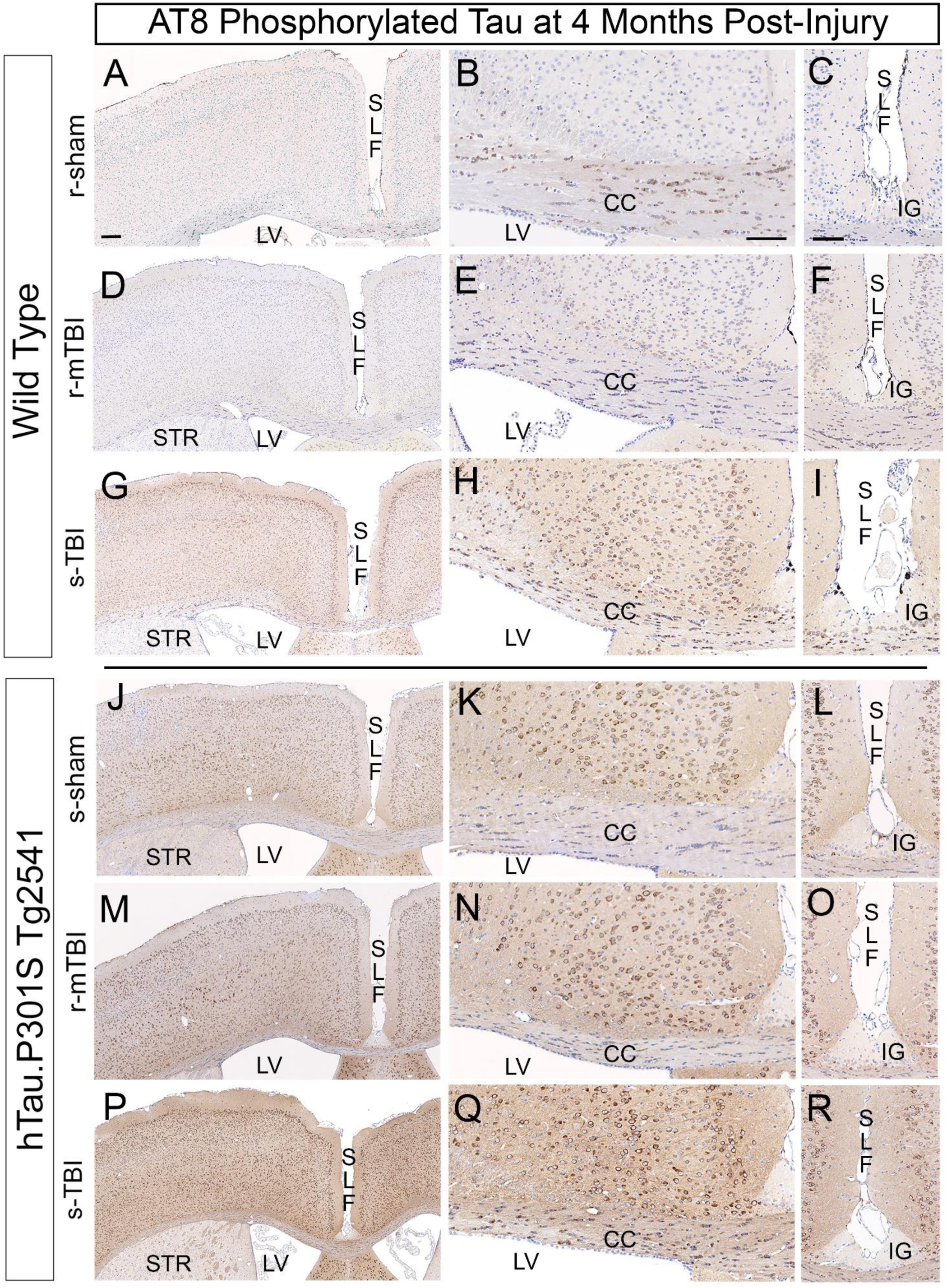
AT8 detection of phosphorylated tau pathology varies with injury conditions and hTau.P301S genotype at 4 months after TBI or sham procedures. A-C. Sham wild type mice do not exhibit AT8 immunolabeling of cortical neurons. In contrast, strong AT8 immunolabeling is found in oligodendrocytes in the corpus callosum (CC). D-F. Wild type r-mTBI mice exhibit AT8 in the cytoplasm of oligodendrocytes, which typically appear as small round cells that are aligned in rows within the CC and sparsely distributed in cortical regions. G-I. Wild type s-TBI mice exhibit clear AT8 immunolabeling in cortical neurons, particularly in superficial layers, along with CC thinning. J-K. In homozygous sham mice, AT8 immunolabels cortical neurons and callosal oligodendrocytes. M-O. Homozygous r-mTBI mice exhibit cortical AT8 immunolabeling, along with CC thinning. P-R. Homozygous s-TBI mice exhibit strong cortical AT8 immunolabeling, along with CC thinning. A-R. Nuclei stained blue with hematoxylin. Scale bars: left column shown in A = 200 µm; middle column in B = 100 µm, right column in C = 100 µm. Abbreviations: Superior longitudinal fissure (SLF), lateral ventricle (LV), corpus callosum (CC), striatum (STR), indusium griseum (IG).

**Table 2.** Localization of phosphorylated tau pathology in coronal sections under the impact site

| Injury | hTau | Corpus callosum |  | Deep CTX layers IV-VI | Sites relevant to chronic traumatic encephalopathy |  |  |  |  |  |
| --- | --- | --- | --- | --- | --- | --- | --- | --- | --- | --- |
|  |  |  |  |  | superficial CTX |  | SL fissure depth |  | CTX peri-vascular cells | peri-ventricular |
|  |  | cells | axons |  | pial or subpial (I) | layers II/III | peri-vascular | layer I |  |  |
| s-sham | WT | ++ | - | + | + | + | +/- | + | - | - |
| **s-TBI** | WT | ++ | + | ++ | ++ | +++ | + | + | +/- | + |
| r-sham | WT | ++ | - | +/- | + | + | + | +/- | - | - |
| **r-mTBI** | WT | + | - | +/- | + | + | ++ | - | +/- | +/- |
| s-sham | Hom | + | + | ++ | + | ++ | + | + | + | + |
| **s-TBI** | Hom | ++ | ++ | +++ | +++ | ++++ | + | + | + | + |
| r-sham | Hom | + | + | +++ | +++ | +++ | + | ++ | +/- | + |
| **r-mTBI** | Hom | + | + | +++ | +++ | +++ | ++ | + | + | + |

CTX = cerebral cortex; SL = superior longitudinal; Roman numerals designate layers of cerebral cortex
Note: Thy1.2 promoter in Thy1-YFP-16 mice labels neurons in cortical layers II-VI (Feng et al., 2000).

In wild type mice, coronal brain sections show increasing AT8 immunolabeling in cells under the impact site with increasing injury severity (Figure 3A-I). This increase was most evident in the medial cerebral cortex, particularly in the superficial cortical layers, after the moderate s-TBI (Figure 3G). Sham mice exhibit little AT8 in cortical neurons while strong AT8 immunolabeling was present in the cytoplasm of cells with the morphology of oligodendrocytes in the corpus callosum (Figure 3B).

In homozygous mice, strong AT8 immunolabeling was found in cortical neurons in sham and injured mice (Figure 3J-R). The intensity of AT8 immunoreactivity in the medial cortex appeared to increase with injury severity to be most pronounced after the s-TBI (Figure 3P). After s-TBI, AT8 immunolabeling of neuronal processes was also notable across cortical layers and in the corpus callosum (Figure 3Q).

The superior longitudinal fissure region was also examined as potentially analogous to human cortical sulci, which are sites of AT8 accumulation in cases of CTE [19] [45]. In wild type and homozygous mice, labeled cells were present along the pial surface in the fissure (Figure 3). Rare AT8 immunolabeled cells were also present in the depth of the fissure along the vasculature or the junction of the gray matter over the corpus callosum (Figure 3L, O, R). Interestingly, the region of the indusium griseum is not immunolabeled with AT8, which supports the specificity of the AT8 signal in cortical neuron processes in the adjacent tissue (Figure 3L, O, R). The indusium griseum is a band of non-cortical cells that runs rostro-caudally superior to the corpus callosum in human brain [13] and rostral mouse brain [1], which may limit comparison of the rostral superior longitudinal fissure to human cortical sulci.

### Endogenous mouse tau is phosphorylated in cortical neurons during chronic phase TBI in wild type mice

Further analysis of AT8 immunolabeling confirmed endogenous mouse tau phosphorylation in cortical neurons of wild type mice at 4 months after s-TBI (Figure 4A-D). HT7 immunolabeling is specific for human tau and does not recognize endogenous mouse tau. After s-TBI in wild type mice, cortical neurons were clearly immunolabeled with AT8, in the absence of HT7 immunolabeling (Figure 4A-B). In contrast, after s-TBI in homozygous mice, cortical neurons were strongly immunolabeled for AT8 with corresponding HT7 immunolabeling (Figure 4C-D). HT7 immunostaining was present in neurons across cortical layers 2-6 (Figure 4D), as expected from the Thy1.2 neuronal promoter [23].

**Figure 4.**
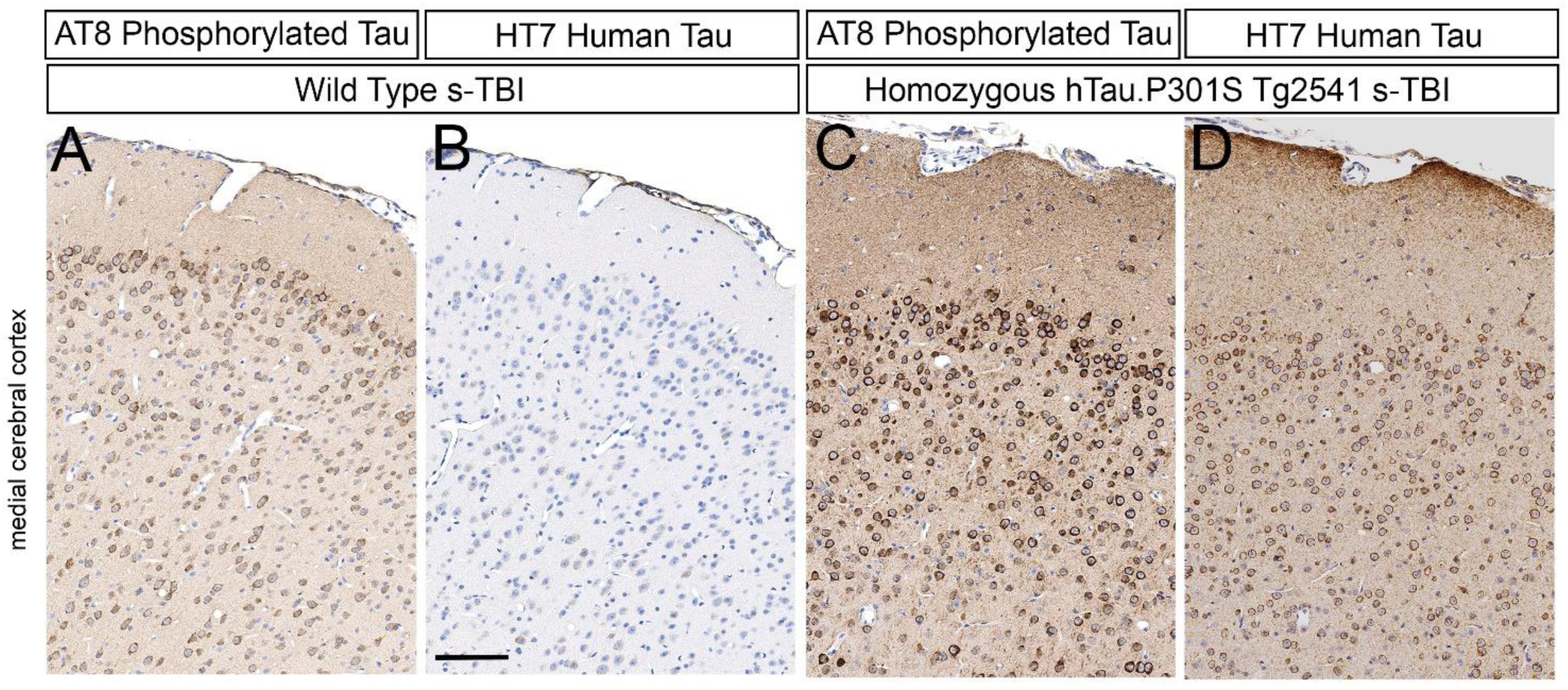
Demonstration of endogenous mouse tau phosphorylation in cortical neurons at 4 months after s-TBI in comparison with human tau expression. A-B. After s-TBI in wild type mice, cortical neurons exhibit clear AT8 immunolabeling of phosphorylated endogenous mouse tau, which is not labeled with the HT7 antibody. C-D. After s-TBI in homozygous mice, cortical neurons exhibit strong AT8 immunolabeling and HT7 detection of human tau. A-D. Nuclei stained blue with hematoxylin. Scale bars for A-D shown in B = 200 µm.

### Phosphorylated Tau is localized in axons during chronic phase TBI in tau homozygous mice

In addition to AT8 immunolabeling of cortical neuron cell bodies, axons were also clearly immunolabeled with AT8 in injured homozygous mice at 4 months after s-TBI (Figure 5). Cortical neurons of the medial cerebral cortex (anterior cingulate cortex, motor cortex) project axons through the corpus callosum that descend in bundles within the striatum [17] [25]. In agreement with this pathway, AT8 immunolabeling of axons was evident in fibers within the corpus callosum (Figure 5A, B) and descending through the striatum (Figure 5E, F) in Tg2541 homozygous s-TBI mice. Striatal neurons lacked AT8 immunolabeling (Figure 5A, E), even in the presence of human tau transgene expression (Figure 5B, F). AT8 immunoreactivity was not evident in axons of the corpus callosum (Figure 5C, D) or striatum (Figure 5G, H) in absence of injury in wild type mice.

**Figure 5.**
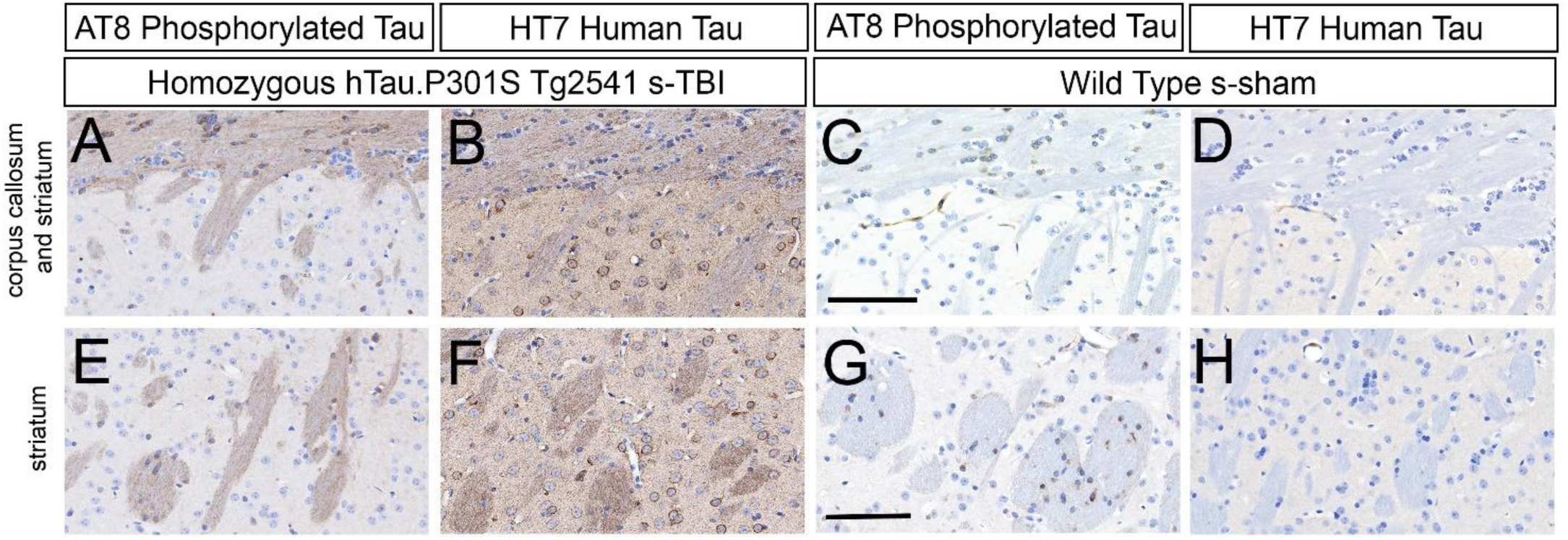
Tau phosphorylation in axons and oligodendrocytes. A-B. After s-TBI in homozygous mice, AT8 immunolabels axons and small round oligodendrocytes in the CC. Striatal neurons underlying the CC are not immunolabeled with AT8, but do express the human transgene as detected with the HT7 antibody. C-D. In sham wild type mice, AT8 labels oligodendrocytes in the CC and underlying striatum. A lack of HT7 signal confirms that the oligodendrocyte AT8 signal reflects endogenous mouse tau phosphorylation. E-F. After s-TBI in homozygous mice, deeper striatal regions show AT8 immunolabeling of cortical neuron axon bundles and small round oligodendrocytes. The neurons are not immunolabeled with AT8, but do express the human transgene as detected with HT7. G-H. In sham wild type mice, deeper striatal regions show distinct AT8 immunolabeling of oligodendrocytes near axon bundles, in the absence of HT7 immunostaining. A-H. Nuclei stained blue with hematoxylin. Scale bars: A-D shown in C = 100 µm, E-H in G = 100 µm.

Analysis of AT8 in comparison to HT7 in the corpus callosum and striatum also revealed striking AT8 immunolabeling of endogenous mouse phosphorylated tau in the cytoplasm of oligodendrocytes (Figure 5). AT8 immunolabeled oligodendrocytes in wild type mice with only the sham procedure (Figure 5C, G). Therefore, oligodendrocytes did not require injury or mutant tau expression to exhibit tau phosphorylation. Oligodendrocytes were not immunolabeled with HT7 in homozygous mice, indicating that AT8 immunolabeling in oligodendrocytes was detecting endogenous mouse tau (Figure 5B, F). This result is in agreement with reports of cytoplasmic tau expression in oligodendrocytes in normal adult mice [48] [77] and after TBI [37].

### Multiple phospho-tau epitopes are increased in brain lysates of tau transgenic mice

Quantitative analysis of brain lysates demonstrated significant increases of total human mutant tau and specific phosphorylated tau epitopes in Tg2541 transgenic mice (Figure 6). Brain lysates were prepared from bilateral regions under the impact site (Figure 6A). This dissection captures corpus callosum axons along with the corresponding callosal projection neurons [89] to account for potential phosphorylation dependent redistribution of tau from the axon resulting in mis-localization in the somatodendritic compartment [39]. These brain regions under the impact site have shown axon damage after s-TBI and r-mTBI in Thy1-YFP-16 mice in our prior studies [53] [88]. Soluble proteins were isolated and probed for tau epitopes using Wes Protein Simple capillary electrophoresis immunoassays (Figure 6B). Soluble tau was quantified as an early stage of phosphorylated tau that is relevant to the axonal compartment and the response to traumatic injury. Soluble tau can disrupt axonal transport and have detrimental effects independent of plaques and tangles [18, 68].

**Figure 6.**
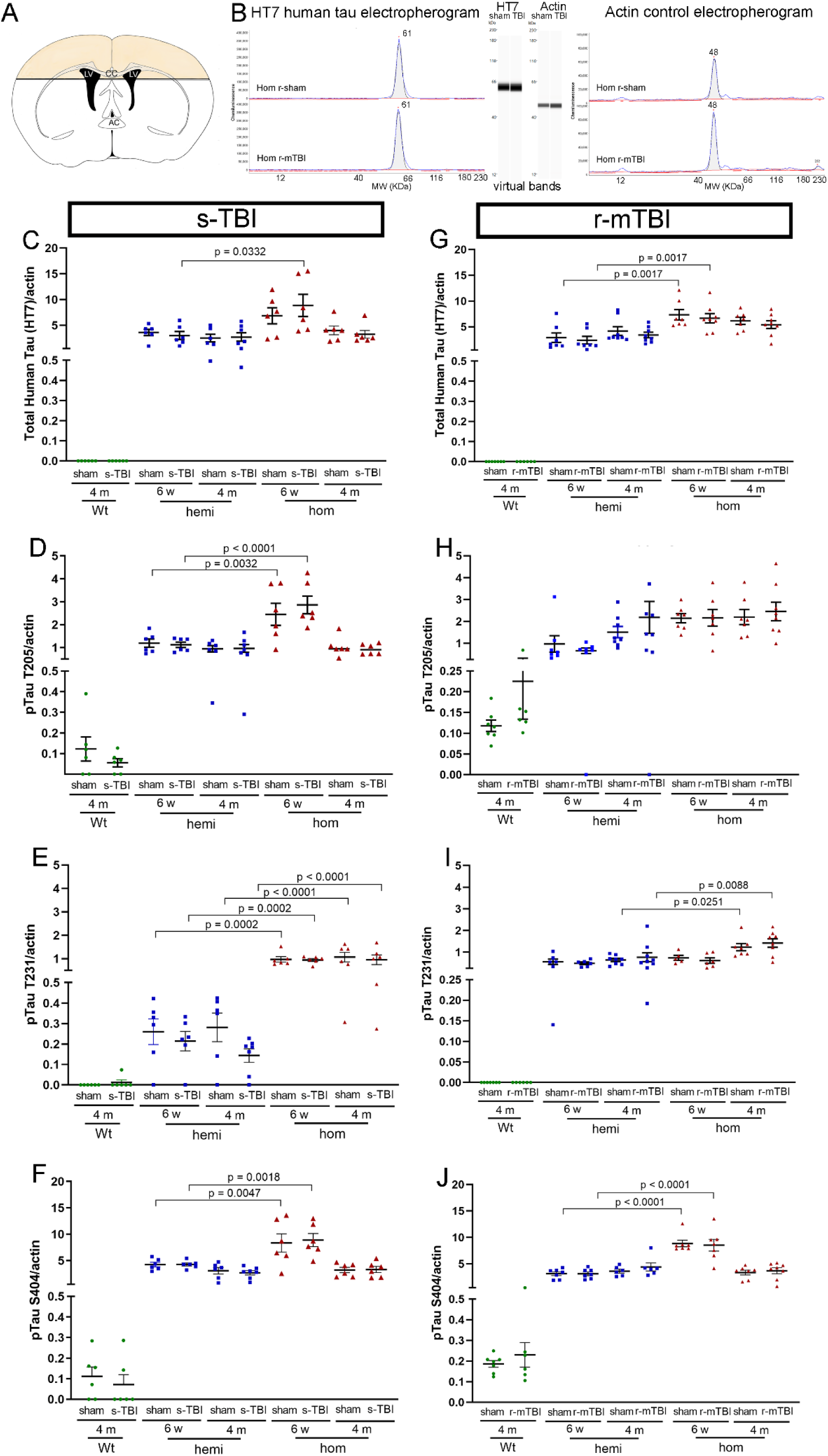
Total human tau and phosphorylated tau are increased in brain lysates of tau transgenic mice. A. Brain lysates were prepared from the colored cortical regions and white matter tracts, including the corpus callosum (CC) over the lateral ventricles (LV). This superior brain region was dissected from a 2-mm thick coronal slice to capture cortical neuron cell bodies and corresponding axons. B. Tau epitopes were quantified in the soluble fraction from brain lysates using Protein Simple Wes capillary electrophoresis immunoassay with actin for normalization of the protein loading amounts. The automated quantification is displayed as electropherograms with the same data also displayed as virtual bands in a blot-like image. C-J. The HT7 antibody recognizes total human tau to detect the human transgene expression. In contrast, the antibodies to phosphorylated tau epitopes recognize both human and mouse epitopes. Dots represent individual mice with wild type (Wt) in green, hemizygous (hemi) in blue, and homozygous (hom) in red. Non-significant comparisons are not shown. Full statistical analysis provided in Table SI-5. C-F. Comparison across genotypes for s-TBI and s-sham mice at 6 weeks and 4 months post-injury. G-J. Comparison across genotypes for r-mTBI and r-sham mice at 6 weeks and 4 months post-injury.

Soluble human tau protein was recognized with the HT7 antibody in wild type, hemizygous, and homozygous Tg2541 mice after s-TBI (Figure 6C-F) or r-mTBI (Figure 6G-J). HT7 did not detect wild type mouse tau when run with the same protein loading and antibody dilution as the transgenic mice (Figure 6C, G). Phosphorylated tau epitopes (pT205, pT231, pS404) were markedly increased in transgenic mice. Wild type mice also exhibited phosphorylation of endogenous mouse tau for T205, which is one of two tau epitopes recognized by AT8, and for S404, which is associated with axonal tau. Across genotypes, soluble tau and phospho-tau levels were not altered by s-TBI or r-mTBI, as compared to the respective sham procedures in mice of each genotype.

### Chronic white matter pathology is significantly worsened in tau homozygous mice after TBI

In contrast to prior mouse TBI studies [5] [27] [56], we found significant effects of human tau transgene expression on the progression of white matter pathology after TBI (Figures 7-10). Corpus callosum width and myelination measures were used as indicators of post-traumatic neurodegeneration after s-TBI and r-mTBI. Mice of each tau genotype were compared at 6 weeks and 4 months post-injury to assess the progression of white matter pathology at subacute and chronic stages, respectively.

**Figure 7.**
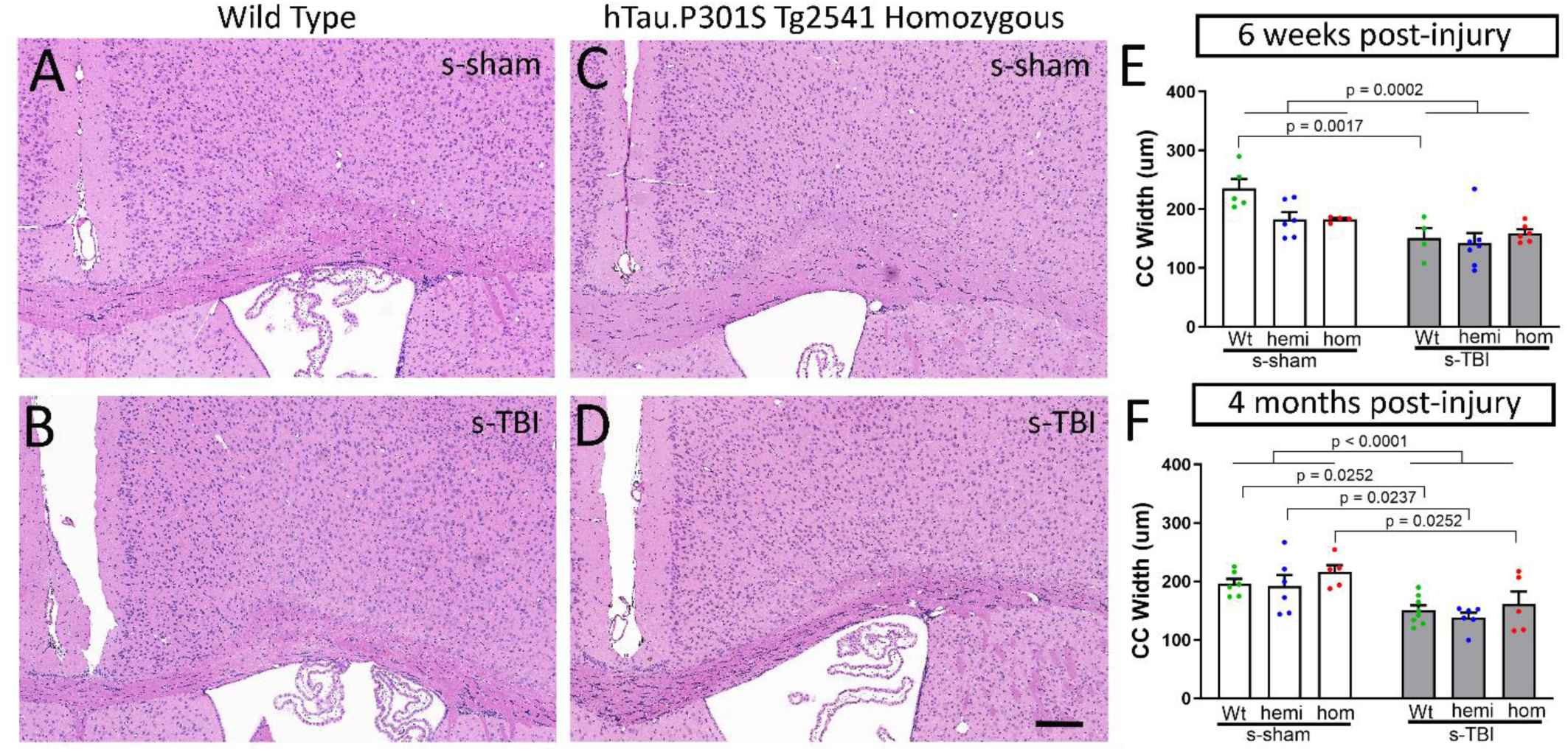
Moderate s-TBI produces corpus callosum atrophy that is not altered by tau genotype. A-D. Coronal sections through the medial cortex and corpus callosum under the impact site with cell nuclei stained with hematoxylin (blue) and cytoplasm with eosin (pink). Corpus callosum thinning is evident at 4 months post-injury after s-TBI, as compared to sham, in wild type (A-B) and tau homozygous mice (C-D). Scale bars for A-D shown in D = 200 µm. E. Corpus callosum width is reduced due to a main effect of injury at the subacute phase. F. Corpus callosum width is significantly reduced due to injury in s-TBI versus sham conditions for each genotype at the chronic phase. Tau genotype does not worsen corpus callosum thinning. E-F. Dots represent individual mice with wild type (Wt) in green, hemizygous (hemi) in blue, and homozygous (hom) in red. Non-significant comparisons are not shown. Full statistical analysis provided in Table SI-3, 4.

A striking difference was observed for the effect of tau genotype on corpus callosum width between injury models. Coronal sections of regions under the impact site were examined with hematoxylin and eosin histological stain, which is widely used for clinical and experimental neuropathology (Figures 7-8). After s-TBI, the corpus callosum width was significantly reduced at both 6 weeks and 4 months post-injury, as compared to sham controls (Figure 7E, F). Tau genotype did not have a significant effect on the corpus callosum width after s-TBI (Figure 7E, F). In contrast, the r-mTBI model did not reduce corpus callosum width at 6 weeks post-injury, as compared to the sham procedure, in mice of any of the three genotypes (Figure 8E). At 4 months post-injury, the corpus callosum thinning was still not observed in wild type or hemizygous mice (Figure 8F). Importantly, significant worsening of corpus callosum thinning at 4 months after r-mTBI in homozygous mice (Figure 8F). Therefore, the repetitive mild closed head injury in mice with high tau pathology resulted in a delayed progression of white matter injury that produced dramatic corpus callosum atrophy.

**Figure 8.**
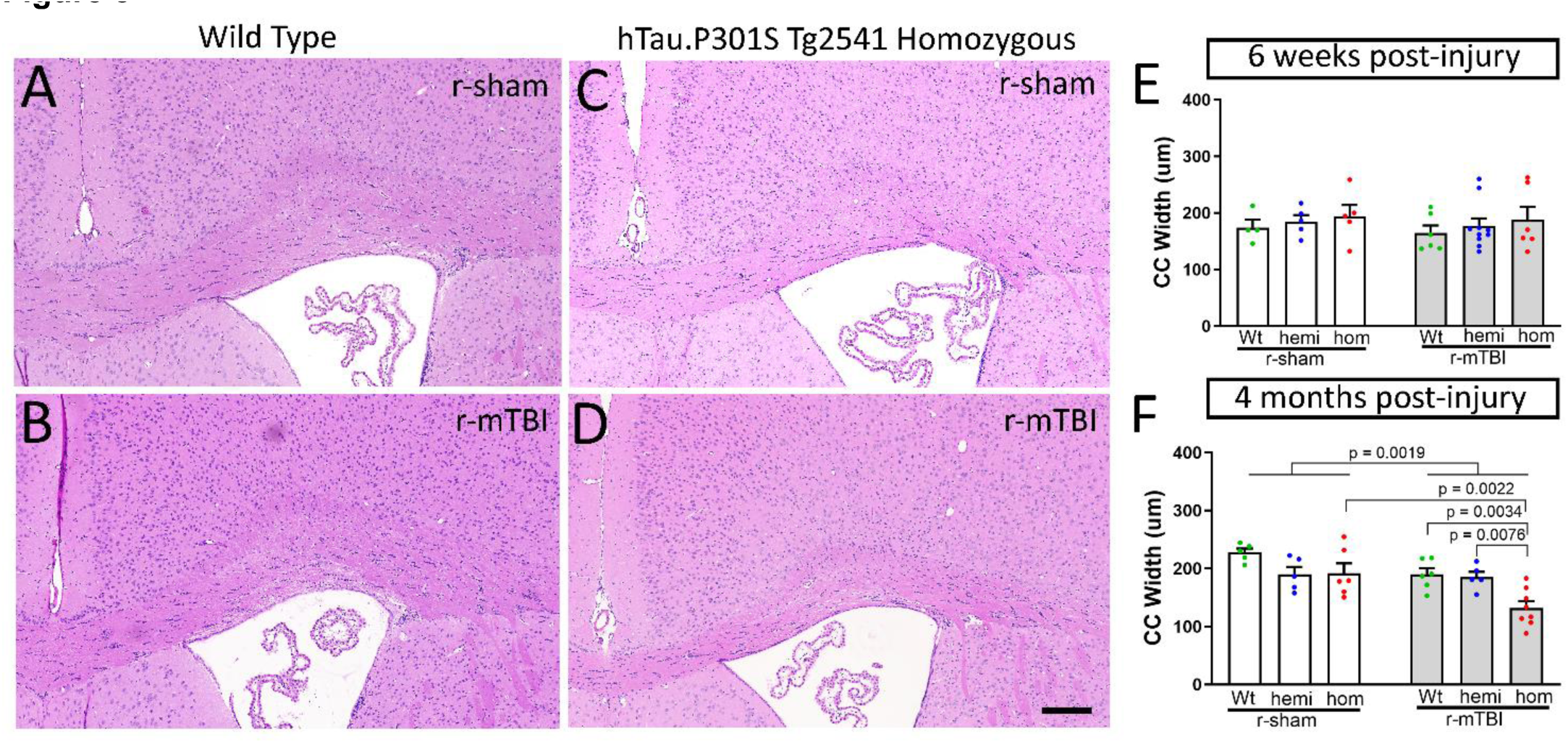
Increased pathological tau produces delayed corpus callosum atrophy in chronic r-mTBI. A-D. Coronal sections through the medial cortex and corpus callosum under the impact site with cell nuclei stained with hematoxylin (blue) and cytoplasm with eosin (pink). Normal adult corpus callosum thickness is seen in wild type mice, both sham (A) and injured (B), and in sham tau homozygous mice (C) at the 4 month time point. Corpus callosum thinning is evident at 4 months post-injury after r-mTBI only in homozygous tau mice (D). Scale bars for A-D shown in D = 200 µm. E. Corpus callosum width is not altered by r-mTBI or tau genotype at the subacute phase. F. The r-mTBI results in a dramatic reduction of corpus callosum width by the chronic phase only when combined with the homozygous tau genotype, but not in wild type and hemizygous mice. E-F. Dots represent individual mice with wild type (Wt) in green, hemizygous (hemi) in blue, and homozygous (hom) in red. Non-significant comparisons are not shown. Full statistical analysis provided in Table SI-3, 4.

Analysis of myelination, with immunolabeling for myelin basic protein (MBP), corroborated the chronic stage findings from the histological stain and revealed an additional effect of tau genotype (Figures 9-10). The delayed corpus callosum thinning after r-mTBI was found only in tau homozygous mice (Figure 9E), matching the findings shown in Figure 8F. The r-mTBI mice did not show significant corpus callosum demyelination, i.e. loss of myelin staining, across genotypes (Figure 9F). After s-TBI, the injury had a significant effect of reduced corpus callosum width that was similar across genotypes (Figure 10E), as in Figure 7F. In addition to this persistent corpus callosum atrophy after s-TBI, significant demyelination developed between the subacute and chronic time points only in the tau homozygous mice (Figure 10F). These results indicate that homozygous mutant tau transgene expression worsened the progression of white matter pathology in both r-mTBI and s-TBI models. The comparison of Wt and hom mice in each injury model demonstrates a large effect size of increased tau in addition to the significant p value determination. The chronic stage onset of corpus callosum atrophy after r-mTBI has a Cohen’s d = 2.16 and the additional pathology of demyelination with atrophy after s-TBI has a Cohen’s d = 2.91, indicating that the hom are more than 98% below the Wt values for each finding.

**Figure 9.**
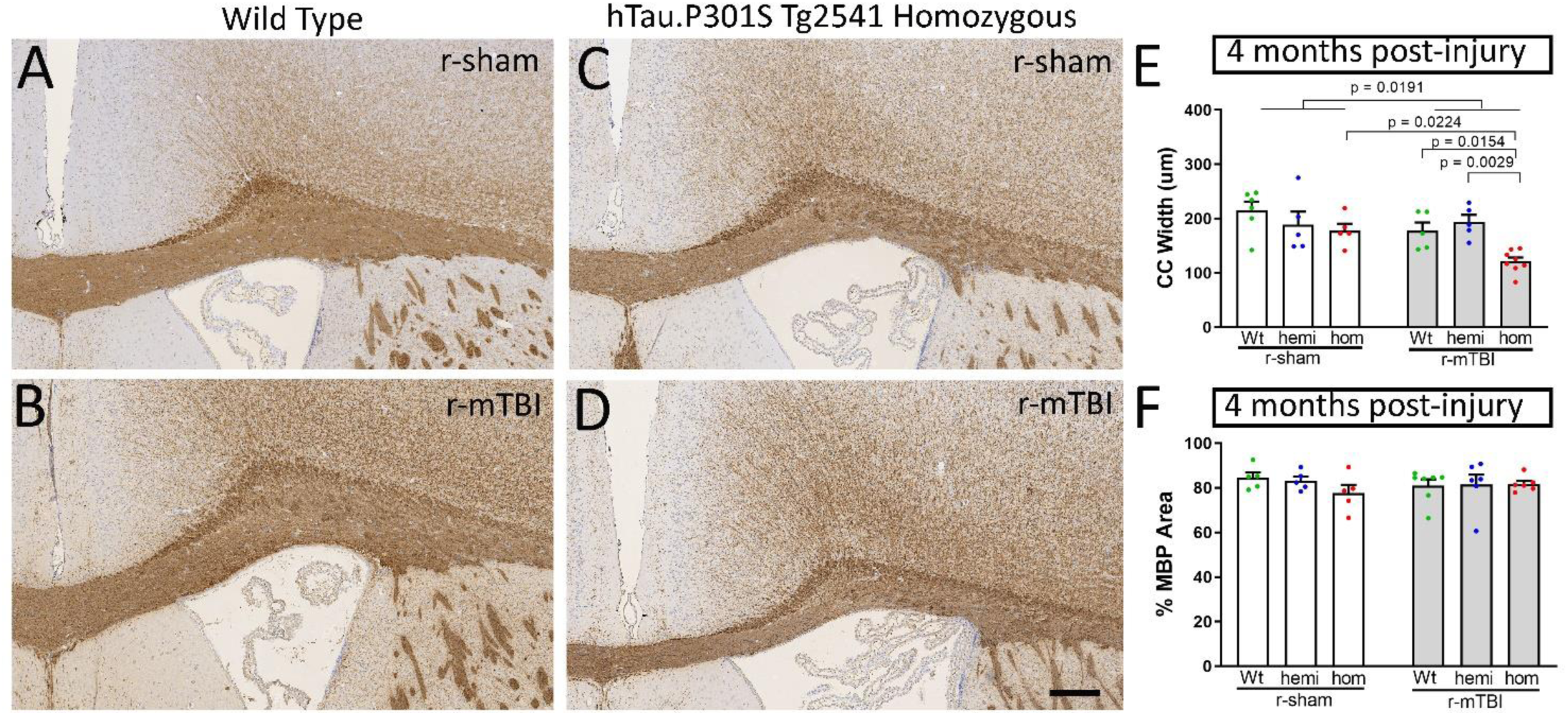
Myelin immunolabeling corroborates delayed corpus callosum atrophy with increased pathological tau in chronic r-mTBI. A-D. Coronal sections through the medial cortex and corpus callosum under the impact site with myelin labeled by immunohistochemistry for myelin basic protein (MBP). Nuclei stained blue with hematoxylin. Normal adult corpus callosum thickness is seen in wild type mice, both sham (A) and injured (B), and in sham tau homozygous mice (C) at the 4 month time point. Corpus callosum thinning is evident at 4 months post-injury after r-mTBI only in homozygous tau mice (D). Scale bars for A-D shown in D = 200 µm. E. The r-mTBI results in a dramatic reduction of corpus callosum width by the chronic phase only when combined with the homozygous tau genotype, but not in wild type and hemizygous mice. F. The r-mTBI did not cause loss of myelinated areas within the corpus callosum across genotypes. E-F. Dots represent individual mice with wild type (Wt) in green, hemizygous (hemi) in blue, and homozygous (hom) in red. Non-significant comparisons are not shown. Full statistical analysis provided in Table SI-3, 4.

**Figure 10.**
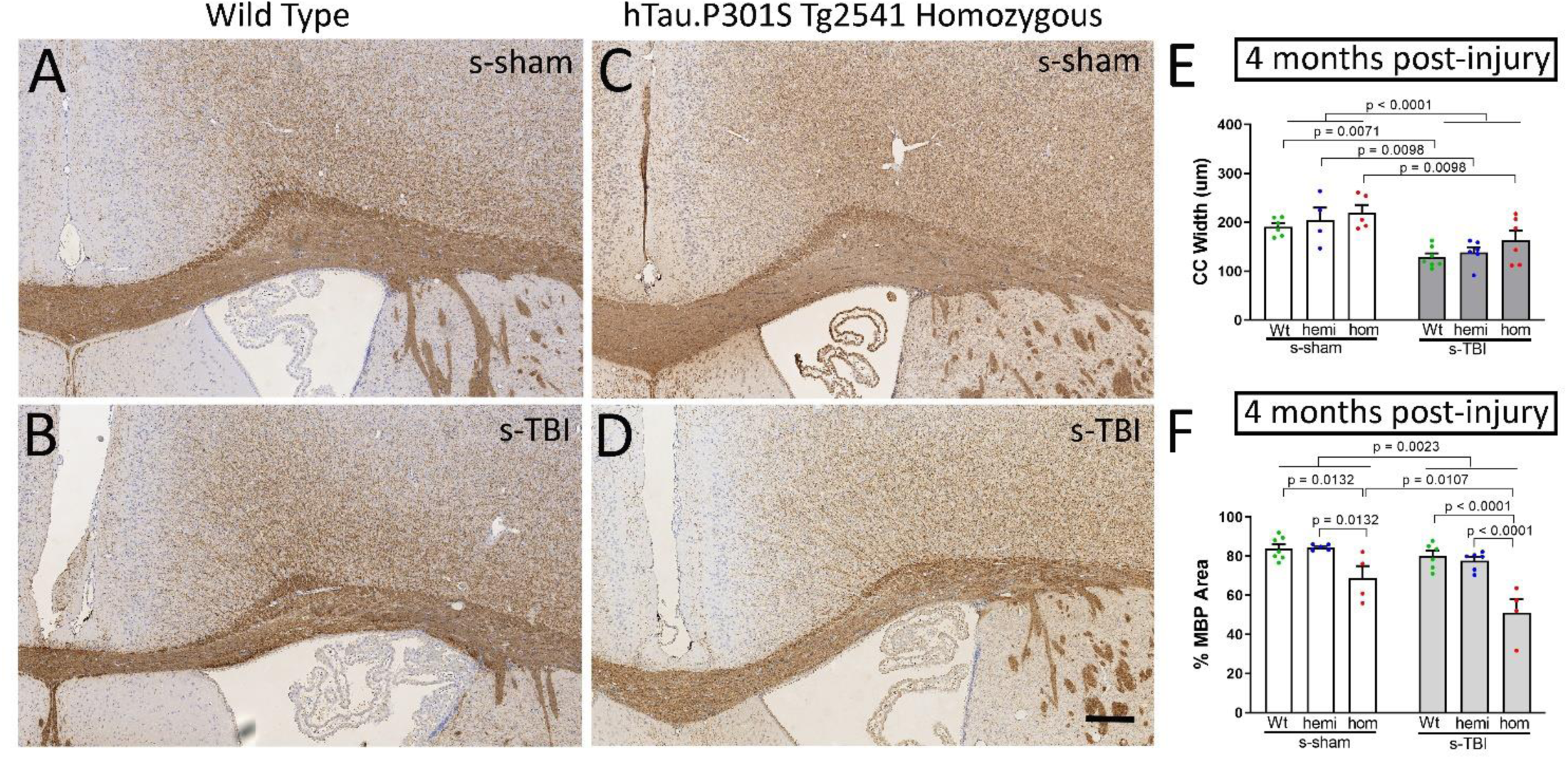
Myelin immunolabeling shows persistent corpus callosum atrophy after s-TBI, and that increased pathological tau produces late stage demyelination. A-D. Coronal sections through the medial cortex and corpus callosum under the impact site with myelin labeled by immunohistochemistry for myelin basic protein (MBP). Nuclei stained blue with hematoxylin. Normal adult corpus callosum thickness is seen in wild type sham mice (A) with thinning after s-TBI (B). Similarly, in tau homozygous mice, the corpus callosum is notably thicker in sham mice (C) as compared to injured mice at the 4 month time point (D). Scale bars for A-D shown in D = 200 µm. E. During chronic phase s-TBI the corpus callosum width is significantly reduced across genotypes. F. Increased tau pathology in homozygous mice results in loss of myelinated areas within the corpus callosum that is significantly worsened in combination with s-TBI. E-F. Dots represent individual mice with wild type (Wt) in green, hemizygous (hemi) in blue, and homozygous (hom) in red. Non-significant comparisons are not shown. Full statistical analysis provided in Table SI-3, 4.

### Both s-TBI and r-mTBI injuries increase microgliosis in the corpus callosum that is enhanced in tau homozygous mice

Neuroinflammation, particularly a chronic microglial response, may contribute to the worsening of chronic stage white matter pathology that was identified with increased tau pathology. Both s-TBI and r-mTBI induced reactive microglia (Figure 11) and astrocytes (Figure 12) in the corpus callosum. This microglial response was significantly increased with tau in homozygous mice after both s-TBI and r-mTBI at the subacute phase and persisted during the chronic phase (Figure 11G-J). The microglial response was more prominent in the white matter than in the adjacent medial cortex under the impact site (Table SI-3 and SI-4). The astrocyte response was not increased in tau homozygous mice (Figure 12G-J). In addition, after r-mTBI, the astrocyte reaction was only significant at the subacute time point and had resolved by the chronic phase (Figure 12J). These results indicate that microgliosis accompanies white matter pathology during the chronic phase post-injury.

**Figure 11.**
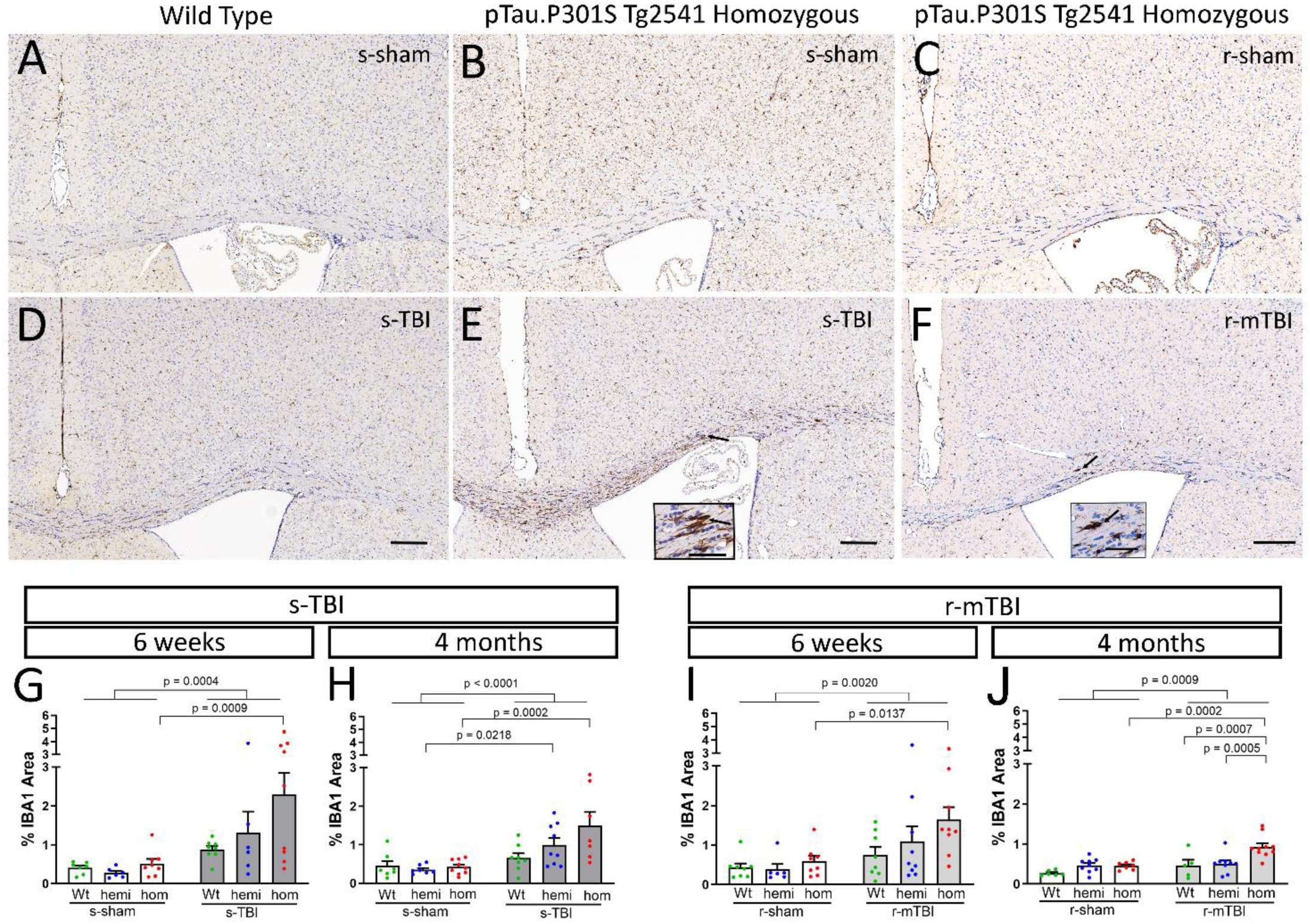
Both s-TBI and r-mTBI injuries induced a microglial response in the corpus callosum that was most prominent in tau homozygous mice. A-F. Immunohistochemistry for IBA1 to identify microglia in coronal sections under the impact site. Nuclei stained blue with hematoxylin. Images shown are 4 months post-injury or sham procedures. Scale bars inset 50 µm, panel images 200 µm. A-C. The microglia appeared similar in the cortex and corpus callosum of sham wild type (A) and tau homozygous mice (B, C). D-F. Both s-TBI (D, E) and r-mTBI (F) appeared to increase IBA1 immunolabeling of microglia in the corpus callosum, with prominent enlargement and elongation after s-TBI in homozygous mice (E). G-J. Quantification of IBA1 immunolabeling in the corpus callosum shows that the tau homozygous genotype is associated with a significant response to injury at both injury models and at both post-injury time points. Dots represent individual mice with wild type (Wt) in green, hemizygous (hemi) in blue, and homozygous (hom) in red. Non-significant comparisons are not shown. Full statistical analysis provided in Table SI-3, 4.

**Figure 12.**
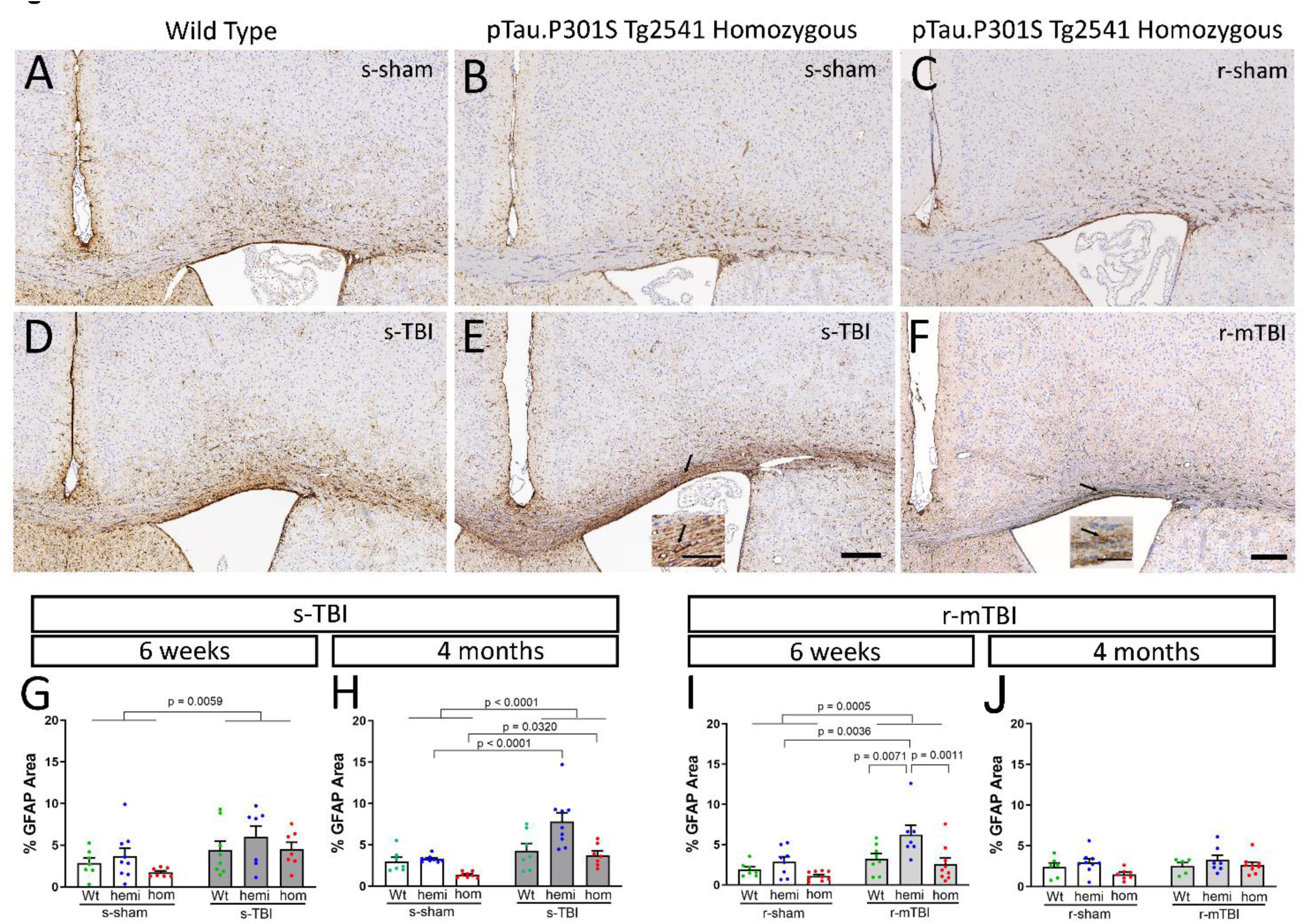
Both s-TBI and r-mTBI injuries induced reactive astrogliosis in the corpus callosum, which resolved in the chronic phase after r-mTBI. A-F. Immunohistochemistry for GFAP to identify astrocytes in coronal sections under the impact site. Nuclei stained blue with hematoxylin. Images shown are 4 months post-injury or sham procedures. Scale bars inset 50 µm, panel images 200 µm. A-C. Astrocyte immunolabeling appeared similar in the cortex and corpus callosum of sham wild type (A) and tau homozygous mice (B, C). D-F. Astrogliosis in the corpus callosum was evident at 4 months after s-TBI (D, E) but localized to a more limited region after r-mTBI (F). G-J. Quantification of GFAP immunolabeling in the corpus callosum shows a main effect of injury at 6 weeks after s-TBI (G) and r-mTBI (I) that persists to 4 months after s-TBI (H) but resolves after r-mTBI (J). The tau homozygous genotype does not significantly increase astrogliosis in either injury models or at either post-injury time point. Dots represent individual mice with wild type (Wt) in green, hemizygous (hemi) in blue, and homozygous (hom) in red. Non-significant comparisons are not shown. Full statistical analysis provided in Table SI-3, 4.

### Translational blood biomarker assay detects axonal injury and human tau exposure levels

A blood biomarker assay used in clinical TBI research [14] [49, 66] was effective as a translational non-invasive method to monitor axonal injury and tau exposure from mouse serum samples (Figure 13). The Simoa® Neurology 4-Plex A was used to test for simultaneous analysis of tau, neurofilament (Nf-L), glial fibrillary acidic protein (GFAP), and ubiquitin carboxyl-terminal hydrolase L1 (UCH-L1) in a given mouse serum sample. This clinical assay effectively detected human tau expressed from the hTau.P301S transgene that continued to be present at high levels in serum of homozygous mice across acute (1 day), subacute (6 weeks), and chronic (4 months) time points (Figure 13A-F). The Neurology 4-Plex A detects mouse Nf-L that is appropriately eliminated with immunodepletion, but is not likely to detect mouse GFAP or mouse UCH-L1 (see Methods), in agreement with our results (data not shown). Nf-L levels were sensitive to axon damage that reflected the injury effect at the acute phase (Figure 13G, J). With progression to the chronic phase, Nf-L levels were dramatically elevated in homozygous mice relative to wild type for s-sham (41.4 fold), s-TBI (45.6 fold), r-sham (32.3 fold) and r-mTBI (49.2 fold) mice. This elevated serum Nf-L level indicates prolonged axon degeneration with continued human mutant tau expression (Figure 13I, L). Both the s-TBI and the r-mTBI produced similar findings for human tau and mouse Nf-L biomarker profiles.

**Figure 13.**
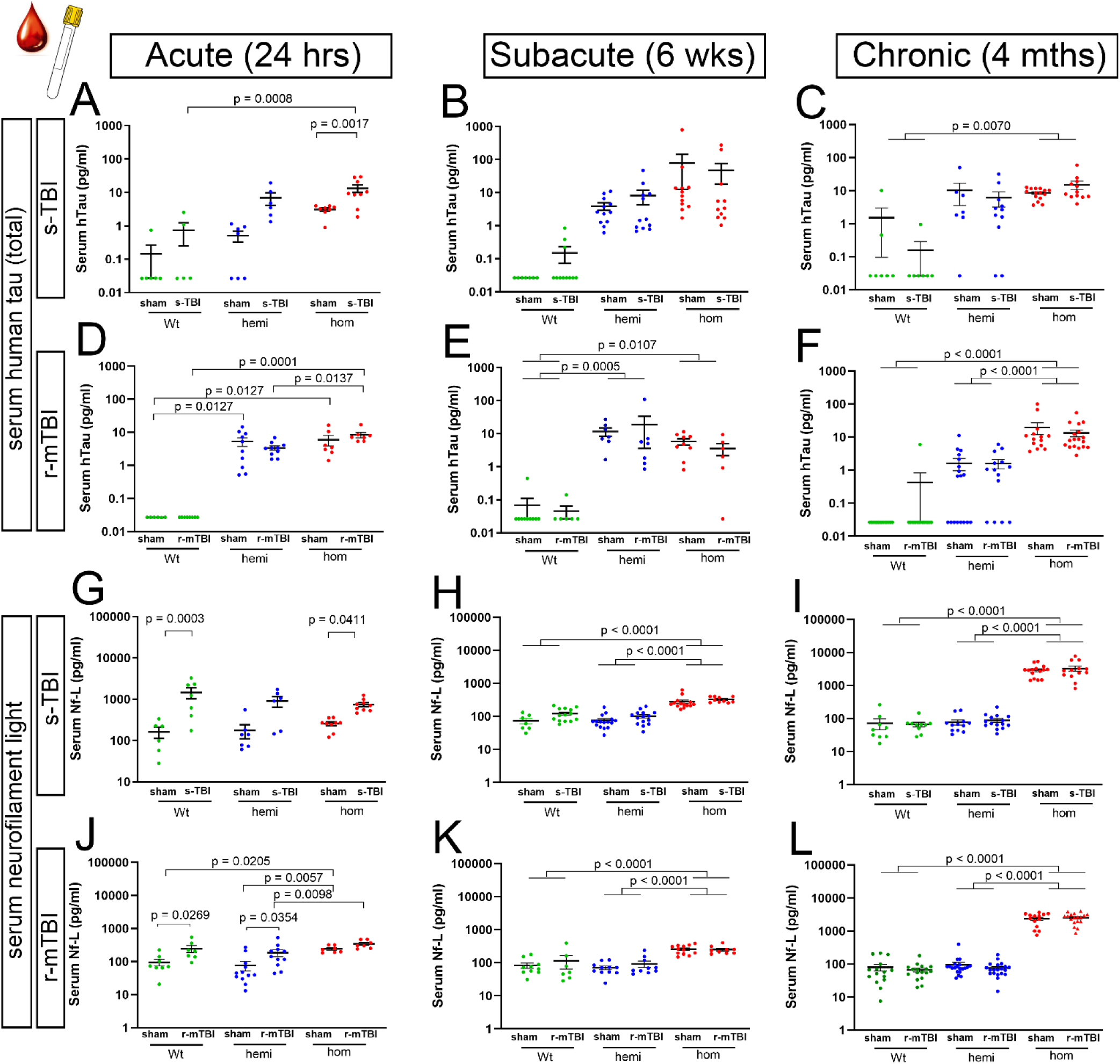
Clinically used blood biomarker assay shows persistent elevation of human mutant tau while mouse neurofilament light detects axonal injury. A-L. Mouse serum analysis using Simoa® Neurology 4-Plex A to simultaneously detect human total tau and mouse neurofilament light (Nf-L) protein levels. Symbols represent individual mice. Non-significant comparisons are not shown. Full details of statistical analysis in Table SI-6. A. In the acute phase, tau protein levels were elevated after s-TBI in serum from homozygous Tg2541 mice, as compared to matched sham mice or to s-TBI wild type mice. B-C. At subacute and chronic time points, human tau protein levels were consistently elevated in s-sham and s-TBI homozygous mice. D-F. Human tau protein levels were consistently elevated in r-sham and r-mTBI mice. In contrast to s-TBI mice, the r-mTBI mice did not show an effect of injury in the acute phase. G, J. In the acute phase, mouse Nf-L protein levels in serum showed a significant injury effect, which was most pronounced after s-TBI in wild type mice. H-I, K-L. Nf-L showed a significant increase in sham and injured homozygous mice that was similar in s-TBI and r-mTBI cohorts. The subacute elevation of Nf-L in homozygous mice further increased at the chronic stage.

## DISCUSSION

The main finding of this study is that increasing pathological tau worsens white matter degeneration after TBI, which is most notable as late phase corpus callosum atrophy after repetitive mild TBI. Demonstrating this interplay over time between tau pathology and long-term outcomes after TBI is challenging in post-mortem human specimens and has remained elusive in animal models. However, a recent study in wild type mice using a closed head injury model similar to our r-mTBI, and supported with a systematic review of the literature in wild type strains, demonstrated the suitability of mouse TBI models to replicate pertinent human tau histopathology [45]. Yet, a recent study and systematic review of TBI in mouse strains expressing human tau constructs failed to identify evidence of a role of tau in neuropathological sequelae after TBI [5] [56].

Our results reveal a complex interplay of increasing pathological human tau relative to the extent of white matter injury and progression to a chronic stage. We bred mice of the hTau.P301S (Tg2541) strain to increase phosphorylated tau pathology in relevant neuron and axon populations to evaluate the effects of 0, 1, or 2 human mutant tau alleles. In the context of a low level of axon damage in the r-mTBI model, increased pathological tau in homozygous mice resulted in significant progression of corpus callosum atrophy between the subacute (6 weeks) to chronic (4 months) time points (Figures 8E-F, 9E). In the moderate s-TBI model, which produces more extensive axonal damage in the corpus callosum, corpus callosum atrophy began in the subacute phase and persisted at chronic time points, regardless of tau genotypes (Figures 7E-F, 10E). Still, the combination of s-TBI with increased tau in homozygous mice worsened pathology of myelinated axons between the subacute and chronic phases based on our findings of late demyelination in the corpus callosum (Figure 10F). Interestingly, the homozygous genotype increased the microglial response to injury at both the subacute and chronic phase in both s-TBI and r-mTBI models (Figure 11).

The P301S human tau mutation was chosen for the current studies as an effective means of increasing phosphorylated tau pathology to inform interpretation of neuropathological sequelae in human TBI cases. These results also support a role of TBI as a modifiable risk factor in frontotemporal dementia (FTD). The human P301S tau mutation, which is expressed in the Tg2541 transgenic mice, is found in early onset FTD [2] [86]. Among FTD patients, suffering a TBI was associated with early onset and sporadic FTD [46] [72]. The interplay of tau pathology and TBI emphasizes the need to identify the direct and/or indirect mechanism(s) by which TBI pathophysiology initiates tau phosphorylation in axons and the subsequent tau pathology, which remain areas of intense research.

The contribution of traumatic axonal injury in white matter atrophy, as demonstrated in our prior studies in *Sarm1* null mice, further supports an interplay of neuronal tau pathology and TBI in our current results. The s-TBI produced more axon damage in the corpus callosum than r-mTBI (Figure 1D-E, SI2), and developed corpus callosum atrophy at an earlier time post-injury. Deletion of the *Sarm1* gene inactivates the SARM1 pathway that executes a highly conserved molecular process of axon degeneration [24] [65]. Acute axonal degeneration at 3 days after s-TBI is dramatically reduced in *Sarm1* null mice as compared to wild type mice [52]. During the chronic phase at 10 weeks after s-TBI, *Sarm1* null mice have both reduced axon degeneration and reduced corpus callosum atrophy [10].

White matter atrophy, demyelination, and neuroinflammation develop after TBI in wild type mice, yet our results show a significant impact of phosphorylated tau pathology on each of these pathological features in the progression from the subacute to chronic phase (Figures 7-12). In the s-TBI model, we have shown persistent axon damage and corpus callosum atrophy by neuropathology and MRI fractional anisotropy with volume analysis at 8 and 10 weeks post-injury [53] [10]. White matter atrophy progressed from proximate to remote sites after a single TBI without detectable phosphorylated tau pathology in neurons [37]. Repetitive mild TBI resulted in corpus callosum atrophy, neuroinflammation and behavioral deficits with longer post-injury intervals of 6 and 12 months [57], as compared to the 4 month time point for our chronic r-mTBI analysis in tau homozygous mice. Additionally, corpus callosum atrophy and reduced fractional anisotropy were found at 6 and 12 months after repetitive mild TBI, and increased plasma neurofilament light levels at one week after repetitive mild TBI correlated with corpus callosum atrophy on MRI at 12 months [55].

Neuropathologically, demyelination is classically identified as a decrease in the area or density of myelin staining [74], as in our finding of reduced myelin staining in the corpus callosum at 4 months after s-TBI months in homozygous mice (Figure 10F). This loss of myelin stain may reflect a loss of myelinated axons and/or loss of the myelin sheath along surviving axons. Our previous studies using electron microscopy for more specific and sensitive analysis of axon and myelin pathology identified a dispersed pattern of demyelination of intact axons within the corpus callosum after s-TBI [53] [54, 75]. Additional axon pathology is associated with disorganization of the molecular components of the node of Ranvier and the flanking paranodes, where myelin sheaths attach to axons. Node of Ranvier abnormalities have been identified in this s-TBI mouse model, in a swine rotational TBI model, and in human TBI cases [53] [61, 71]. Demyelination can lead to further neurodegeneration since myelination protects vulnerable denuded axons and provides bioenergetic and trophic support [64] [50]. Furthermore, myelin is highly enriched in cholesterol, which is emerging with apolipoprotein E as an important factor in white matter impairment in aging and neurodegenerative diseases [6] [8] [67].

Neuroinflammation is a critical factor in white matter degeneration after TBI and in other neurodegenerative diseases [19] [21] [36] [58]. Microglial activation can persist for years after a TBI with continued axon degeneration, which is associated with corpus callosum atrophy [44]. In agreement with these reports, the microglial response in the corpus callosum was exacerbated in tau homozygous mice after s-TBI and r-mTBI (Figure 11), indicating a potential role in the interaction of tau and TBI in white matter atrophy, demyelination, and/or neuroinflammation.

Interpretation of microglial activation in the current results in Tg2541 mice, with hTau.P301S expressed in neurons from the Thy1.2 promoter, may gain from comparison with the PS19 mice, which have hTau.P301S expressed in multiple cell types from the murine prion promoter (Table SI1). In the absence of injury, adult PS19 mice appear to have broad robust microglial activation in the brain [87]. In contrast, in our studies of sham mice, the microglial response was not different across wild type, hemizygous or homozygous Tg2541 genotypes (Figure 11). In agreement with our post-injury results, repetitive closed head injury increased the microglial response in PS19 mice [42]. The PS19 prion promoter may be advantageous in the context of tau spreading, which was accelerated after a moderate-severe penetrating TBI, and resulted in learning and memory impairment [22]. We conducted initial neurobehavioral testing based on social interaction with the 3-chamber assessment that differentiated r-mTBI effects in wild type mice in our prior study [88].

However, the tau genotype in hemizygous Tg2541 mice impaired social interaction in sham mice to a similar extent as r-mTBI in wild type mice (Figure SI-3), indicating that even this lower level of hemizygous mutant tau can obscure neurobehavioral assessments of an injury effect. The current results from analysis of atrophy, neuroinflammation, and demyelination in the corpus callosum white matter are further supported by the effects of P301S hTau expression in a repetitive model (Table SI-1) that involved mainly the optic nerve and tract [16].

There are several limitations that should be considered in the interpretation of these results. The Neurology 4-Plex A was an informative and translational approach with the advantage of simultaneous analysis of human tau and mouse neurofilament light proteins in a given sample (Figure 13). And, while an injury effect was discerned at the acute time point, serum levels are not specific to effects in the brain since human mutant tau is expressed broadly, including in the spinal cord (Figure SI-4). The performance data of the Neurology 4-Plex A for mouse GFAP and UCHL-1 was not available during our initial studies. Future testing may be more sensitive for neurofilament light with a single assay format, while GFAP and UCHL-1 assays would need to be designed specifically for mouse proteins. Another limitation is that mouse models cannot provide a full range of injury parameters that may be important in human TBI. The two closed head TBI mouse models in this study reflect key pathological features of single moderate TBI and repetitive mild TBI as two main forms of human TBI; each model was previously characterized by neuropathology, fluorescent reporter mice, MRI structural and diffusion tensor imaging, and behavioral assessments. In our analysis, the range of neuropathology created by using the two models was advantageous for identifying multiple chronic phase effects of tau. Future studies in other models could capture additional pathological features, such as traumatic vascular injury and gyrencephalic structure.

## Conclusion

To our knowledge, this study is the first demonstration that modifying neuronal tau worsens late stage TBI neuropathological sequelae in the corpus callosum, a major white matter tract associated with clinical evaluation of post-traumatic neurodegeneration. Our results show a significant impact of phosphorylated tau pathology on corpus callosum atrophy, demyelination, and neuroinflammation in the progression from the subacute to chronic phase TBI. The results reveal an interplay of the tau modification and the extent of TBI pathology over the post-injury time course.

These findings support the applicability of mouse models for identifying potential therapeutics to mitigate neuropathological sequelae of TBI. Large scale phosphoproteomic network analysis in this Tg2541 mouse strain revealed protein co-expression modules associated with tau-induced pathologies, with specific tau phosphorylation sites related to the neuroinflammatory response, mitochondrial bioenergetic processes, cholesterol biosynthesis, and postsynaptic density [79].

These modules and networks are also relevant to TBI pathologies. Therefore, future studies leveraging the current results could provide further insight into factors leading to post-traumatic neurodegeneration and potential targets for screening novel therapeutics.

## Acknowledgements

We are grateful to Dr. Tony Wyss-Coray (Stanford University, California, USA) for providing mice to initiate our hTau.P301S Tg2541 colony. We appreciate assistance from core resources including the Preclinical Behavior and Models Core, Biomarkers Core, and Translational Therapeutics Core of the Center for Neuroscience and Regenerative Medicine and The American Genome Center. We thank Drs. Kryslaine Radomski and Genevieve Sullivan for helpful comments on the manuscript.

## Funding

United States Defense Health Agency Program in Brain Injury and Disease Prevention, Treatment, and Research (HU0001-22-2-0002).

## Disclaimer Statement

The authors have no conflicts of interest to disclose. The views, information or content, and conclusions presented do not necessarily represent the official position or policy of, nor should any official endorsement be inferred on the part of, the Uniformed Services University, the Department of Defense, the U.S. Government or the Henry M. Jackson Foundation for the Advancement of Military Medicine, Inc.

## Author Contributions

The study concept was developed and designed by Regina Armstrong and Daniel Perl. Experiments were designed by Fengshan Yu and Regina Armstrong. Material preparation, data collection and analysis were performed by Fengshan Yu, Diego Iacono, Chen Lai, Jessica Gill, Tuan Le, Patricia Lee, Gauthaman Sukumar, and Regina Armstrong. The first draft of the manuscript was written by Fengshan Yu and Regina Armstrong and all authors commented on previous versions of the manuscript. All authors read and approved the final manuscript.

## Supplemental Information (SI)

**Supplemental Information Table SI-1:** Selected relevant studies of TBI in human tau mouse lines. Recent systematic reviews provide a more comprehensive discussion of tau pathology after TBI in wildtype and transgenic mice (Bachstetter et al., 2020; Kahriman et al., 2021).

Supplemental Information (SI) Supplemental Information Table SI-1: Selected relevant studies of TBI in human tau mouse lines.
| human tau mouse line<br>(source citation) | Tau protein | promoter | genotype control mice | TBI model (age at time of TBI) | post-injury time points | sex | TBI study citation |
| --- | --- | --- | --- | --- | --- | --- | --- |
| **hTau.P301S (Tg2541)**<br>B6-Tg(Thy1-MAPT*P301S)2541<br>(Allen et al., 2002; Johnson et al., 2017) | human P301S mutation in 0N4R hTau isoform (hemizygous and homozygous) | murine Thy1.2 (neuronal) | wild type littermates | moderate **CHI**<br>1x<br>(8 wks) | 1 dpi, 6 wpi, and 4 mpi | ♂ ♀ | Current Study |
|  |  |  | wild type littermates | mild **r-CHI**<br>1/dy x 5 dys<br>(8 wks) | 1 dpi, 6 wpi, and 4 mpi | ♂ ♀ | Current Study |
| **hTau58.4**<br>(Yin et al., 2017) | human P301S mutation in 0N4R hTau isoform (homozygous) | murine Thy1.2 (neuronal) | C57BL/6 | mild **r-CHI**<br>6/dy x 7 dys<br>(12-16 wks) | 1 and 3 mpi | ♂ | Cheng et al., 2020 |
| **hTau.P301S (PS19)**<br>B6;C3-Tg(Prnp-MAPT*P301S)PS19Vle/J<br>(Yoshiyama et al., 2007) | human P301S mutation in 1N4R hTau isoform (homozygous) | murine prion protein | wild type littermates | moderate-severe **CCI**<br>1x<br>(12 wks) | 1 dpi, 1 wpi, 2 and 6 mpi | not stated | Edwards et al., 2020 |
|  |  |  | C57BL/6J | mild **r-CHI**<br>1/dy x 3<br>(35-41 dys) | 40 dpi | ♂ | Izzy et al., 2021 |
| **hTau**<br>B6.CgMapt<sup>tm1(EGFP)Klt</sup> Tg(MAPT)8cPdav/J<br>(Andorfer et al., 2003) | human MAPT 6 hTau isoforms (hemizygous) and murine MAPT knockout (homozygous) | human MAPT | none | mild **r-CHI**<br>1x and 5x over 10 dys<br>(12 wks) | acute, 6 and 12 mpi | ♂ | Mouzon et al., 2019 |
|  |  |  |  | Mild **r-CHI**<br>1/dy x 20<br>(16 wks) | acute, 3 and 12 mpi | ♂ | Gangolli et al., 2019 |

MAPT = microtubule-associated protein tau
CHI = concussive closed head injury; r-CHI = repetitive CHI
CCI = controlled cortical impact onto dura (with craniotomy)
mpi, wpi, or dpi = months, weeks, or days post injury as reported by study

**Supplemental Information Figure SI-1.**
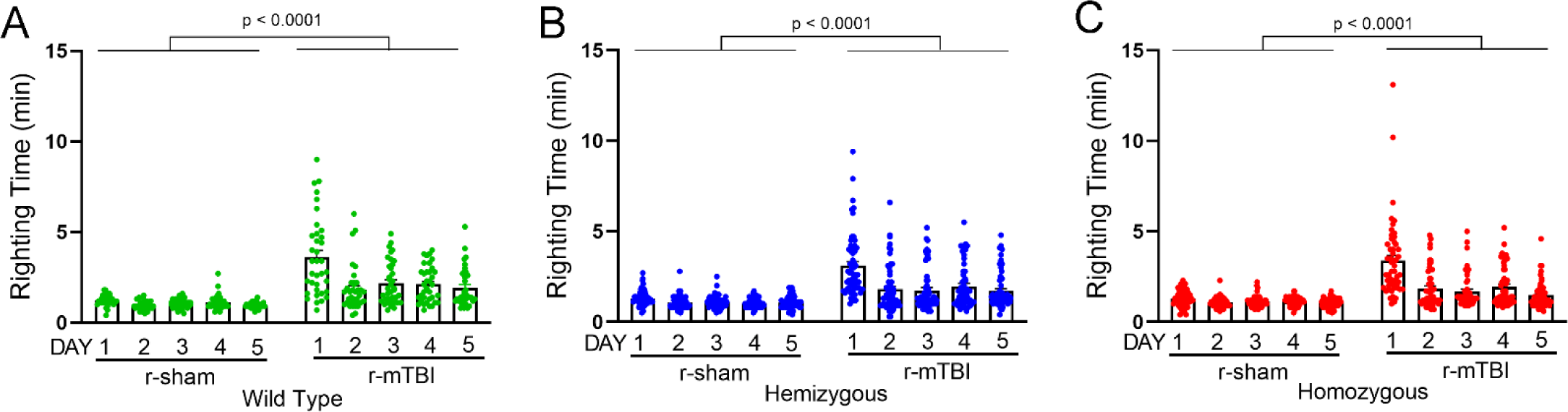
Righting reflex recorded immediately after r-sham or r-mTBI procedures with data shown for all five days for mice of each genotype. See legend to Figure 1.

**Supplemental Information Figure SI-2.**
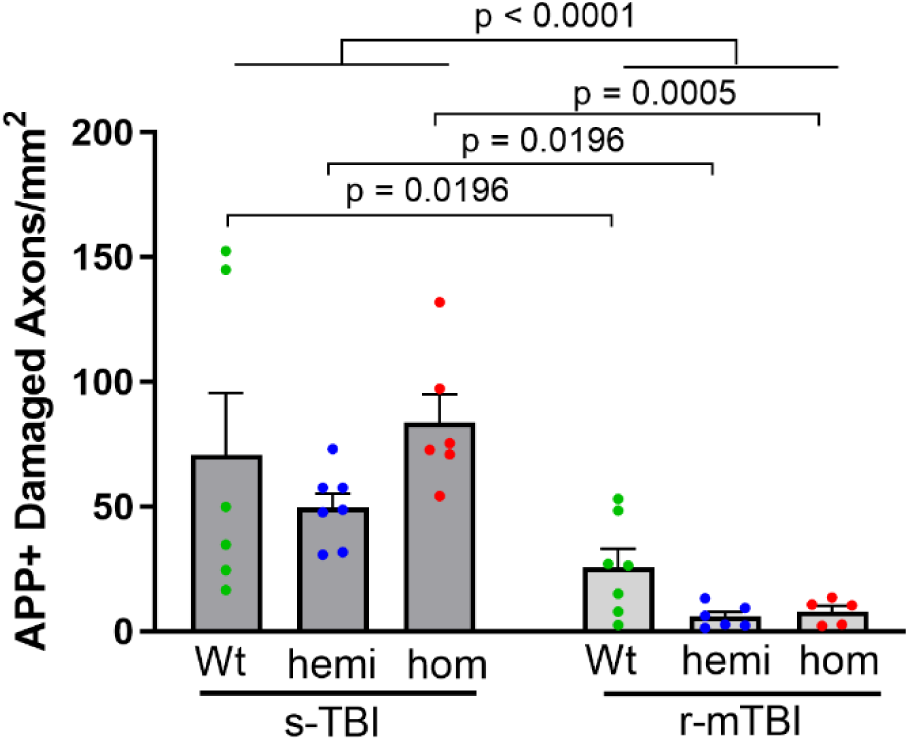
Acute axon damage in the corpus callosum at 24 hours after the moderate single (s-TBI) is more extensive than after the repetitive mild TBI (r-mTBI). See Figure 1.

**Supplemental Information Figure SI-3.** Assessment of social impairment using the 3-chamber assay.

### Methods for social interaction assessment

This r-mTBI model produced social interaction deficits in C57BL/6 adult mice in our prior study (Yu et al., 2017). Therefore, mice of the P301S line were tested for changes associated with TBI or genotype. The social interaction assessment was carried out using a three-chamber sociability test apparatus (Stoelting Co., Wood Dale, IL). The test mouse was first placed in the middle chamber and allowed 10 minutes to explore all chambers for habituation to the apparatus. A small wire enclosure was placed in each of the side chambers and the mouse was allowed to explore all chambers for 10 minutes to habituate to the enclosures as novel objects. Social approach was then tested by placing an unfamiliar C57BL/6J mouse (stranger) of the same sex and age into one of the wire enclosures. The test mouse was again placed into the middle chamber and allowed to explore among the three chambers for 10 minutes. The movement of the test mouse was video recorded and then manually scored to determine the time spent exhibiting social approach behavior (defined as movement toward, circling, or sniffing of the stranger mouse in the wire enclosure) using ANY-Maze software (Stoelting Co.).

**Figure SI-3.**
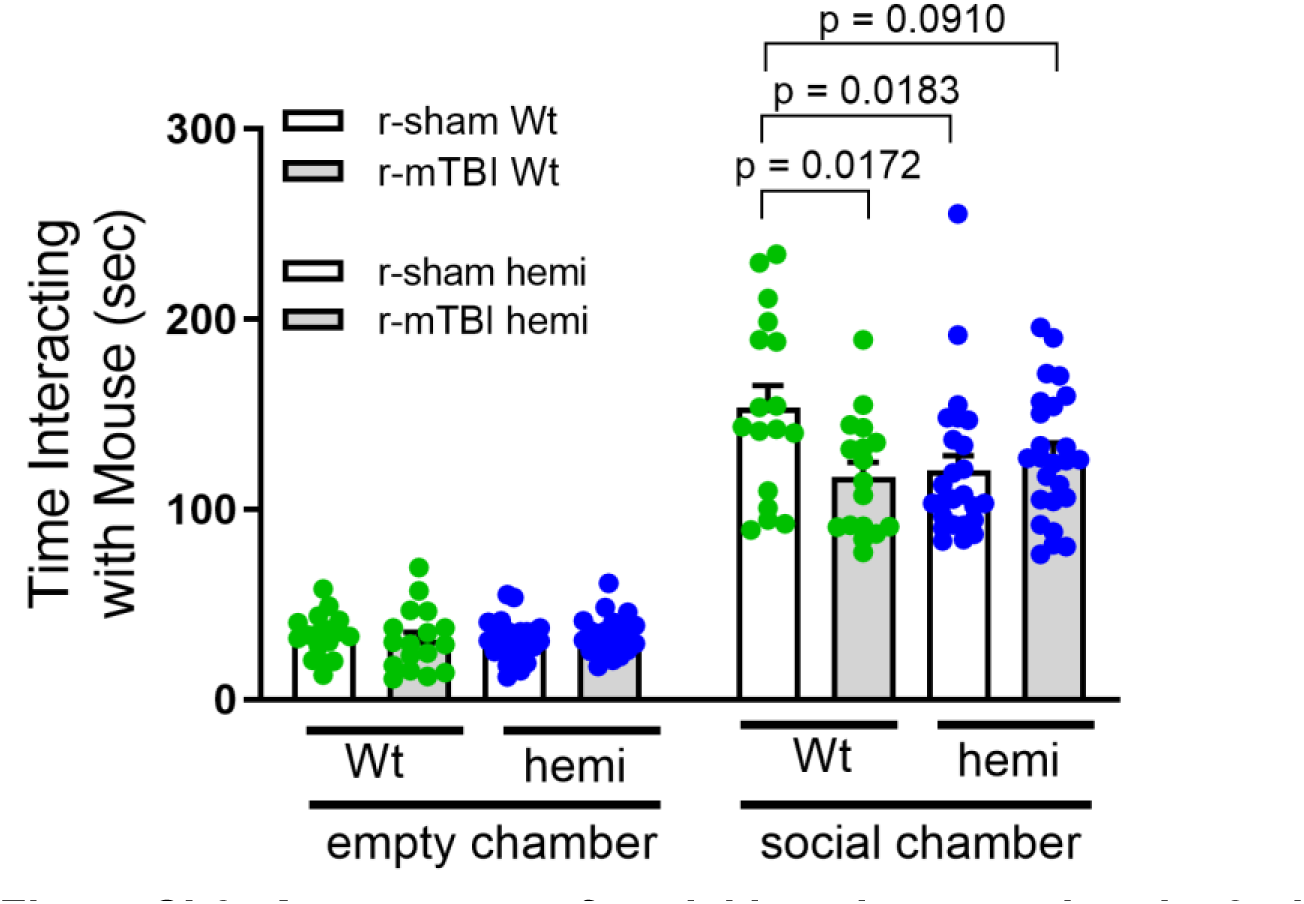
Assessment of social impairment using the 3-chamber assay. After habituation to a central zone, doors are open to two adjacent chambers for the test mouse to choose either the empty chamber with an empty wire carrier or the social chamber with a non-familiar mouse in the wire carrier. Mice of all conditions exhibit a clear preference of entering the social chamber to interact with the non-familiar mouse. The wild type (Wt) injured mice (r-mTBI; Wt, gray bars) spend significantly less time interacting with the non-familiar mouse, as compared to the interaction time for the sham mice (r-sham; Wt, white bars). With hemizygous hTau.P301S Tg2541 mice (hemi), both the r-sham mice exhibit reduced time interacting with the non-familiar mouse, indicating that the tau expression produces social impairment. Male and female mice were tested as separate cohorts. Mice received the injury or sham procedures at 8 weeks of age and were for social testing was conducted 6 weeks later. One-way ANOVA with Dunnett’s test.

**Supplemental Information Figure SI-4.**
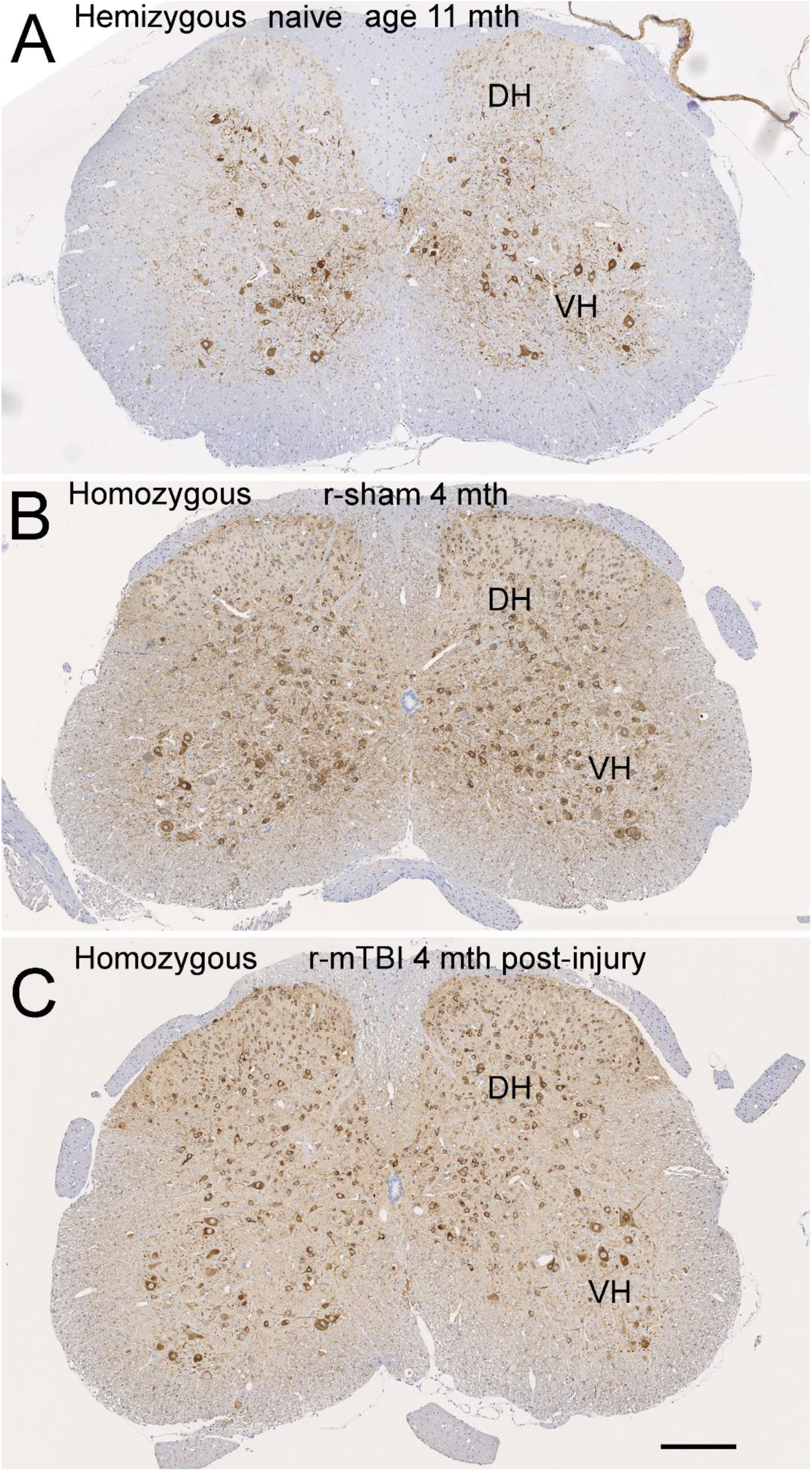
AT8 immunolabeling in spinal cord sections.

**Figure SI-4. *AT8 immunolabeling in spinal cord sections.*** Phosphorylated tau accumulations are seen in sensory neurons of the dorsal horn (DH) and motor neurons of the ventral horn (VH). These examples illustrate tau from regions outside the injury site contribute to total human tau in serum samples. Scale bars = 200 µm.

**Supplemental Information Table SI-2.**
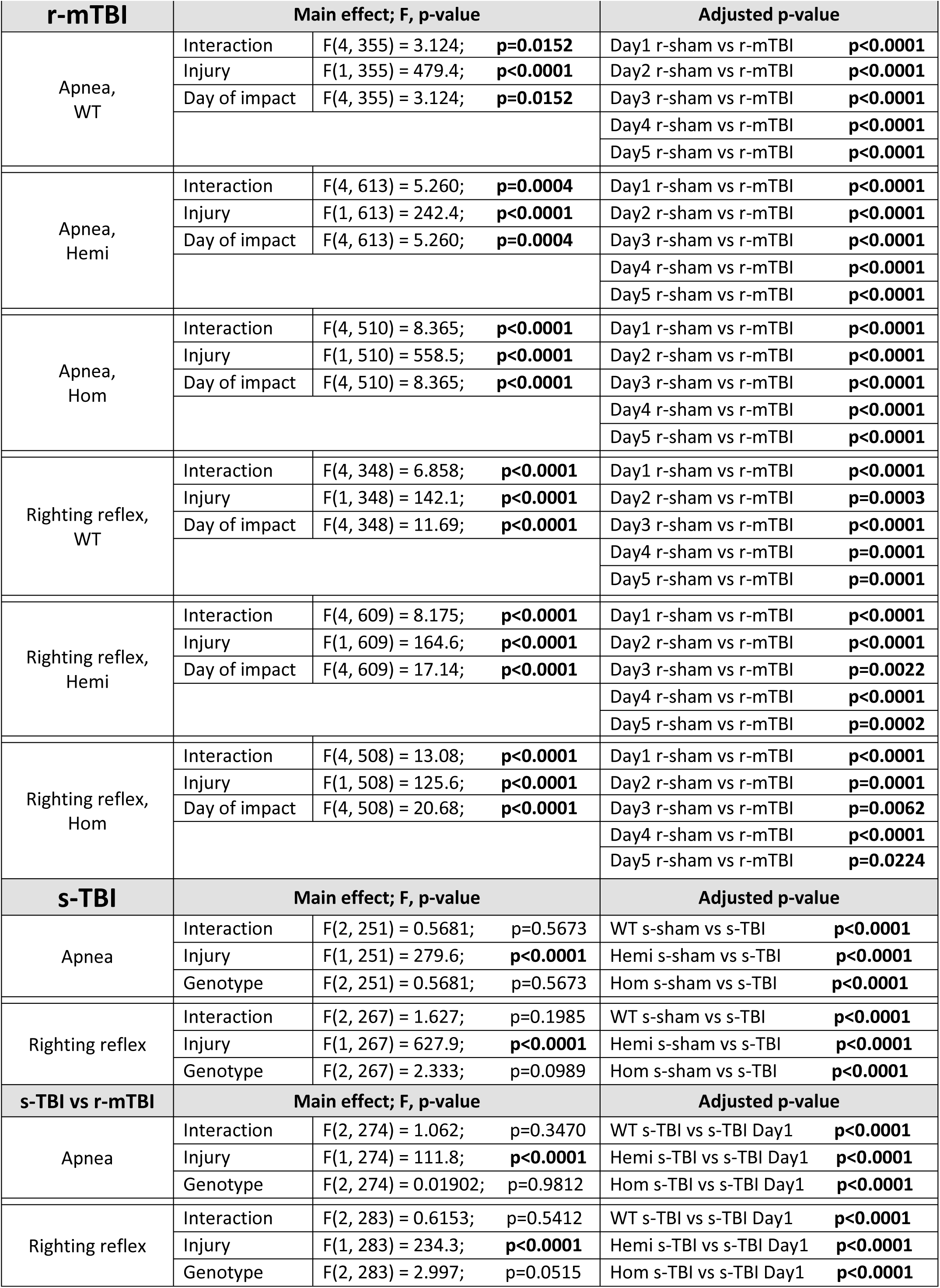
Statistical details for post-surgical assessments.

**Supplemental Information Table SI-3.**
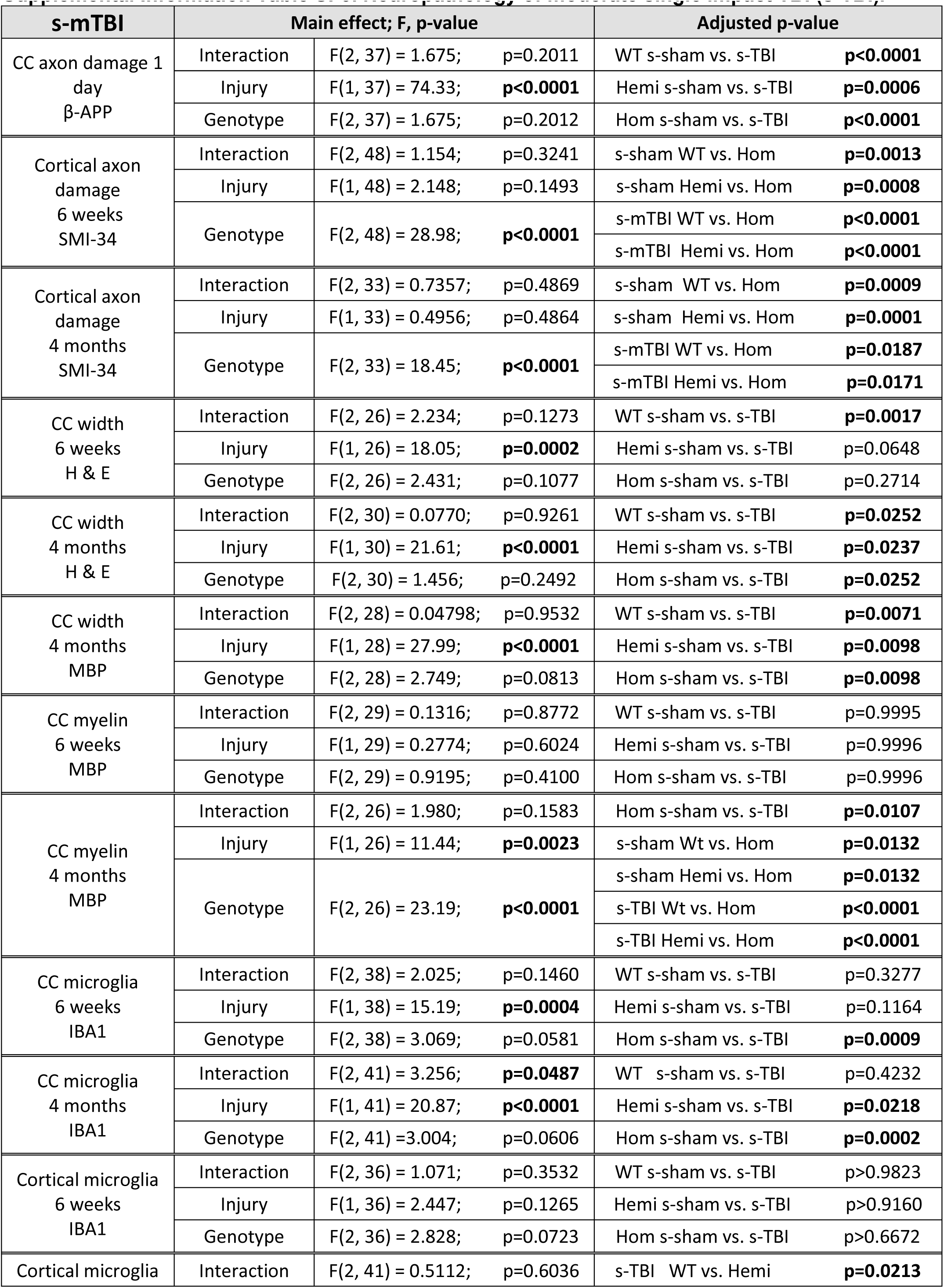
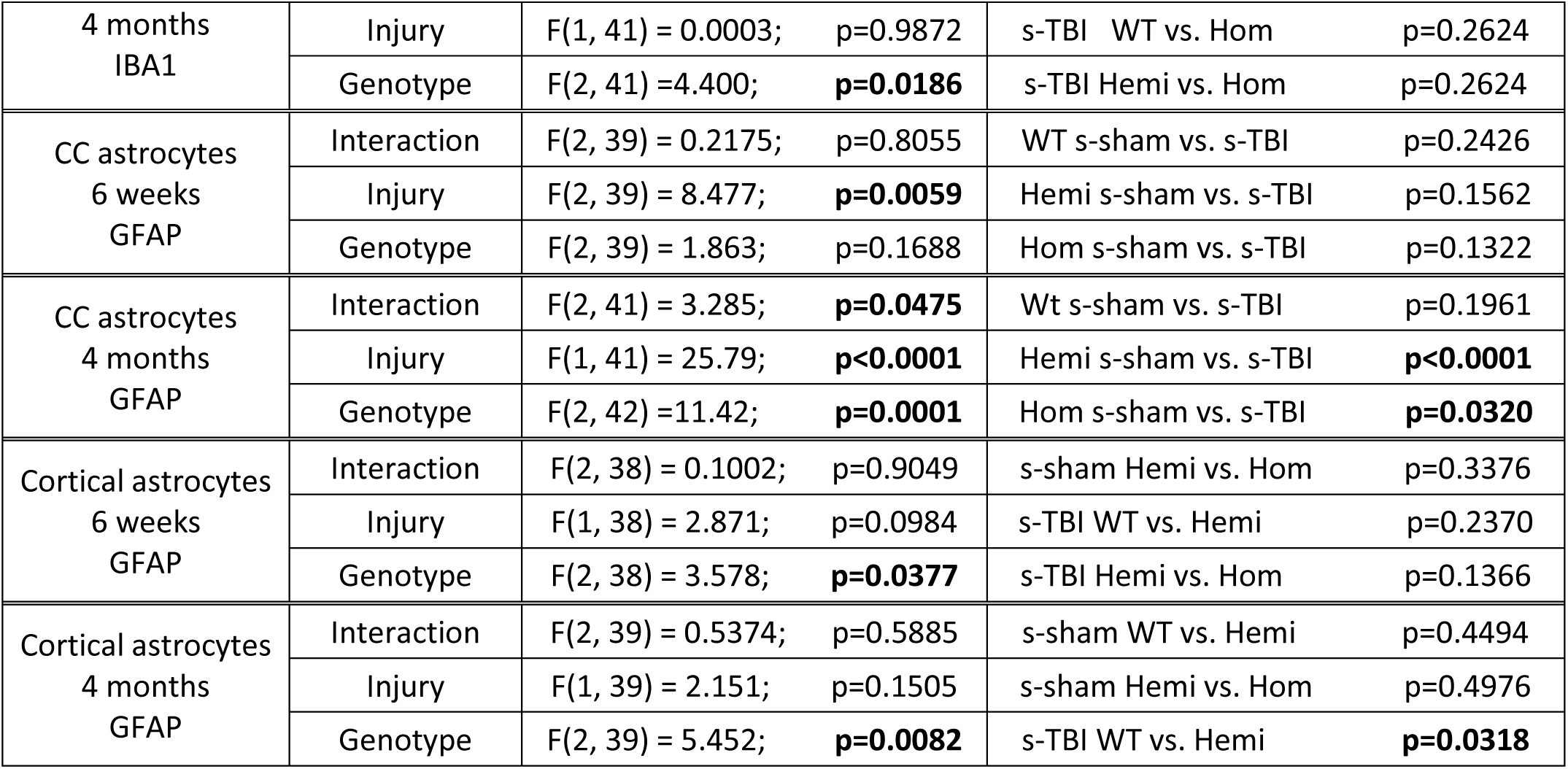
Neuropathology of moderate single impact TBI (s-TBI).

**Supplemental Information Table SI-4.**
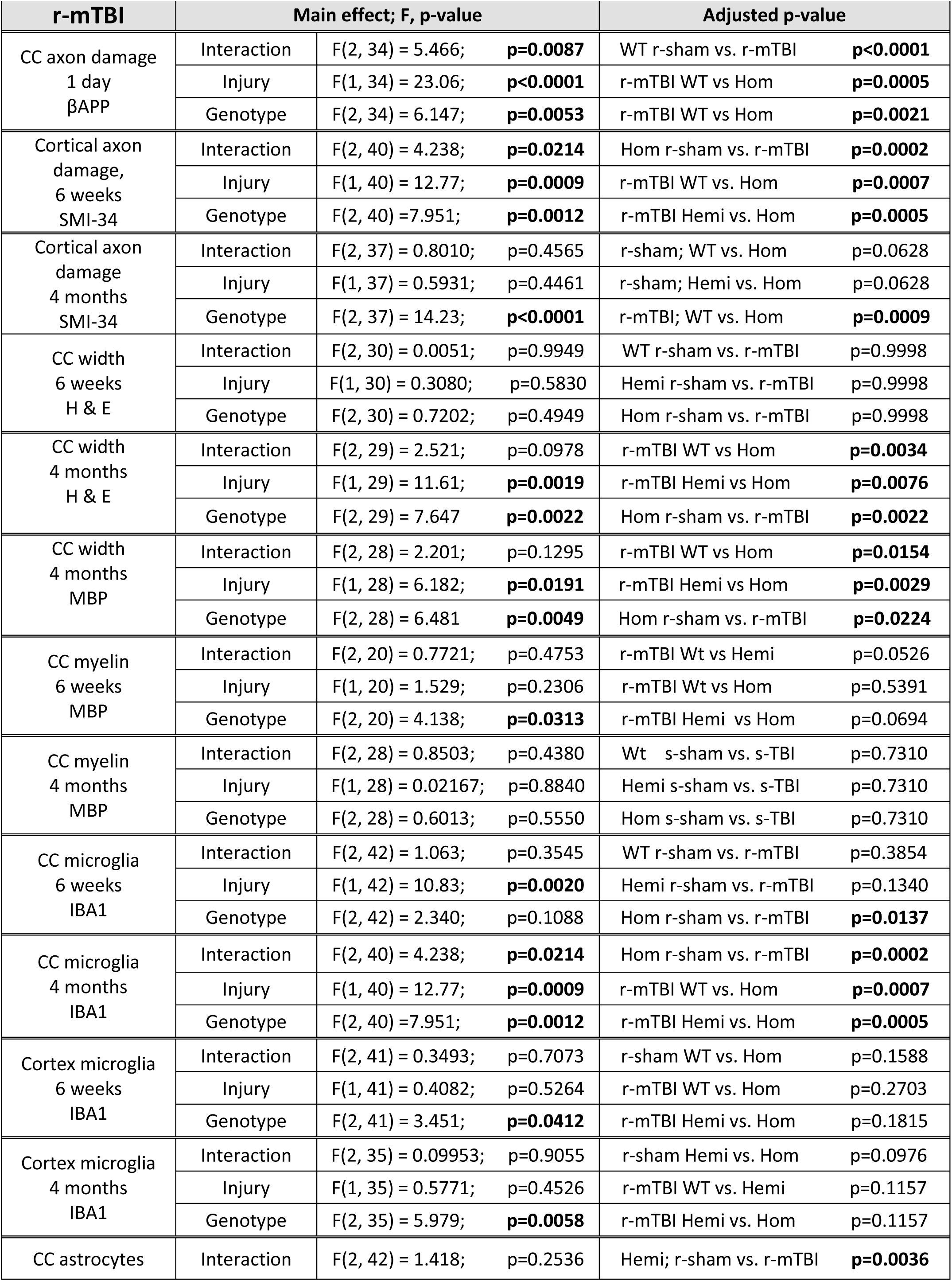
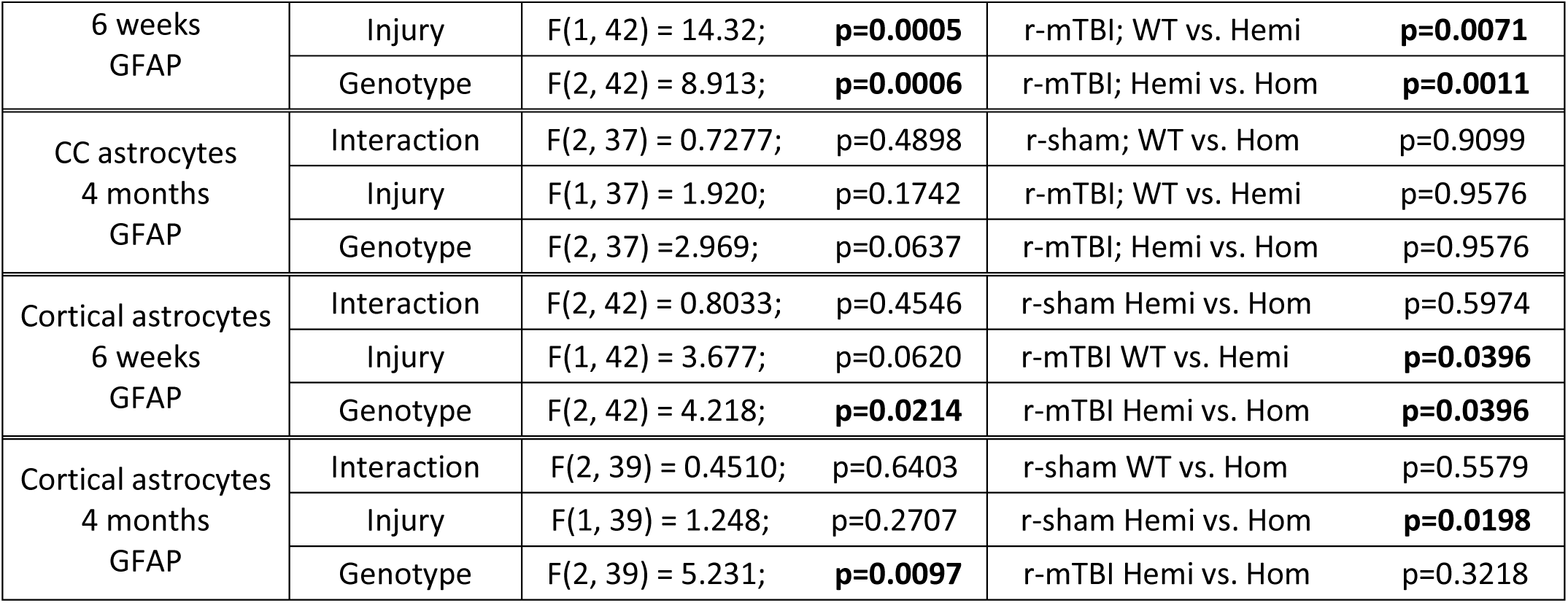
Neuropathology of repetitive mild TBI (r-mTBI).

| **r-mTBI** | **Main effect; F, p-value** |  |  | **Adjusted p-value** |  |
| --- | --- | --- | --- | --- | --- |
| CC axon damage<br>1 day<br>βAPP | Interaction | F(2, 34) = 5.466; **p=0.0087** |  | WT r-sham vs. r-mTBI | **p<0.0001** |
|  | Injury | F(1, 34) = 23.06; **p<0.0001** |  | r-mTBI WT vs Hom | **p=0.0005** |
|  | Genotype | F(2, 34) = 6.147; **p=0.0053** |  | r-mTBI WT vs Hom | **p=0.0021** |
| Cortical axon damage,<br>6 weeks<br>SMI-34 | Interaction | F(2, 40) = 4.238; **p=0.0214** |  | Hom r-sham vs. r-mTBI | **p=0.0002** |
|  | Injury | F(1, 40) = 12.77; **p=0.0009** |  | r-mTBI WT vs. Hom | **p=0.0007** |
|  | Genotype | F(2, 40) = 7.951; **p=0.0012** |  | r-mTBI Hemi vs. Hom | **p=0.0005** |
| Cortical axon damage<br>4 months<br>SMI-34 | Interaction | F(2, 37) = 0.8010; p=0.4565 |  | r-sham; WT vs. Hom | p=0.0628 |
|  | Injury | F(1, 37) = 0.5931; p=0.4461 |  | r-sham; Hemi vs. Hom | p=0.0628 |
|  | Genotype | F(2, 37) = 14.23; **p<0.0001** |  | r-mTBI; WT vs. Hom | **p=0.0009** |
| CC width<br>6 weeks<br>H & E | Interaction | F(2, 30) = 0.0051; p=0.9949 |  | WT r-sham vs. r-mTBI | p=0.9998 |
|  | Injury | F(1, 30) = 0.3080; p=0.5830 |  | Hemi r-sham vs. r-mTBI | p=0.9998 |
|  | Genotype | F(2, 30) = 0.7202; p=0.4949 |  | Hom r-sham vs. r-mTBI | p=0.9998 |
| CC width<br>4 months<br>H & E | Interaction | F(2, 29) = 2.521; p=0.0978 |  | r-mTBI WT vs Hom | **p=0.0034** |
|  | Injury | F(1, 29) = 11.61; **p=0.0019** |  | r-mTBI Hemi vs Hom | **p=0.0076** |
|  | Genotype | F(2, 29) = 7.647 **p=0.0022** |  | Hom r-sham vs. r-mTBI | **p=0.0022** |
| CC width<br>4 months<br>MBP | Interaction | F(2, 28) = 2.201; p=0.1295 |  | r-mTBI WT vs Hom | **p=0.0154** |
|  | Injury | F(1, 28) = 6.182; **p=0.0191** |  | r-mTBI Hemi vs Hom | **p=0.0029** |
|  | Genotype | F(2, 28) = 6.481 **p=0.0049** |  | Hom r-sham vs. r-mTBI | **p=0.0224** |
| CC myelin<br>6 weeks<br>MBP | Interaction | F(2, 20) = 0.7721; p=0.4753 |  | r-mTBI Wt vs Hemi | p=0.0526 |
|  | Injury | F(1, 20) = 1.529; p=0.2306 |  | r-mTBI Wt vs Hom | p=0.5391 |
|  | Genotype | F(2, 20) = 4.138; **p=0.0313** |  | r-mTBI Hemi vs Hom | p=0.0694 |
| CC myelin<br>4 months<br>MBP | Interaction | F(2, 28) = 0.8503; p=0.4380 |  | Wt s-sham vs. s-TBI | p=0.7310 |
|  | Injury | F(1, 28) = 0.02167; p=0.8840 |  | Hemi s-sham vs. s-TBI | p=0.7310 |
|  | Genotype | F(2, 28) = 0.6013; p=0.5550 |  | Hom s-sham vs. s-TBI | p=0.7310 |
| CC microglia<br>6 weeks<br>IBA1 | Interaction | F(2, 42) = 1.063; p=0.3545 |  | WT r-sham vs. r-mTBI | p=0.3854 |
|  | Injury | F(1, 42) = 10.83; **p=0.0020** |  | Hemi r-sham vs. r-mTBI | p=0.1340 |
|  | Genotype | F(2, 42) = 2.340; p=0.1088 |  | Hom r-sham vs. r-mTBI | **p=0.0137** |
| CC microglia<br>4 months<br>IBA1 | Interaction | F(2, 40) = 4.238; **p=0.0214** |  | Hom r-sham vs. r-mTBI | **p=0.0002** |
|  | Injury | F(1, 40) = 12.77; **p=0.0009** |  | r-mTBI WT vs. Hom | **p=0.0007** |
|  | Genotype | F(2, 40) = 7.951; **p=0.0012** |  | r-mTBI Hemi vs. Hom | **p=0.0005** |
| Cortex microglia<br>6 weeks<br>IBA1 | Interaction | F(2, 41) = 0.3493; p=0.7073 |  | r-sham WT vs. Hom | p=0.1588 |
|  | Injury | F(1, 41) = 0.4082; p=0.5264 |  | r-mTBI WT vs. Hom | p=0.2703 |
|  | Genotype | F(2, 41) = 3.451; **p=0.0412** |  | r-mTBI Hemi vs. Hom | p=0.1815 |
| Cortex microglia<br>4 months<br>IBA1 | Interaction | F(2, 35) = 0.09953; p=0.9055 |  | r-sham Hemi vs. Hom | p=0.0976 |
|  | Injury | F(1, 35) = 0.5771; p=0.4526 |  | r-mTBI WT vs. Hemi | p=0.1157 |
|  | Genotype | F(2, 35) = 5.979; **p=0.0058** |  | r-mTBI Hemi vs. Hom | p=0.1157 |
| CC astrocytes | Interaction | F(2, 42) = 1.418; p=0.2536 |  | Hemi; r-sham vs. r-mTBI | **p=0.0036** |

|  |  |  |  |  |
| --- | --- | --- | --- | --- |
| 6 weeks<br>GFAP | Injury | $F(1, 42) = 14.32$; **p=0.0005** | r-mTBI; WT vs. Hemi | **p=0.0071** |
| | Genotype | $F(2, 42) = 8.913$; **p=0.0006** | r-mTBI; Hemi vs. Hom | **p=0.0011** |
| CC astrocytes<br>4 months<br>GFAP | Interaction | $F(2, 37) = 0.7277$; p=0.4898 | r-sham; WT vs. Hom | p=0.9099 |
| | Injury | $F(1, 37) = 1.920$; p=0.1742 | r-mTBI; WT vs. Hemi | p=0.9576 |
| | Genotype | $F(2, 37) = 2.969$; p=0.0637 | r-mTBI; Hemi vs. Hom | p=0.9576 |
| Cortical astrocytes<br>6 weeks<br>GFAP | Interaction | $F(2, 42) = 0.8033$; p=0.4546 | r-sham Hemi vs. Hom | p=0.5974 |
| | Injury | $F(1, 42) = 3.677$; p=0.0620 | r-mTBI WT vs. Hemi | **p=0.0396** |
| | Genotype | $F(2, 42) = 4.218$; **p=0.0214** | r-mTBI Hemi vs. Hom | **p=0.0396** |
| Cortical astrocytes<br>4 months<br>GFAP | Interaction | $F(2, 39) = 0.4510$; p=0.6403 | r-sham WT vs. Hom | p=0.5579 |
| | Injury | $F(1, 39) = 1.248$; p=0.2707 | r-sham Hemi vs. Hom | **p=0.0198** |
| | Genotype | $F(2, 39) = 5.231$; **p=0.0097** | r-mTBI Hemi vs. Hom | p=0.3218 |

**Supplemental Information Table SI-5.**
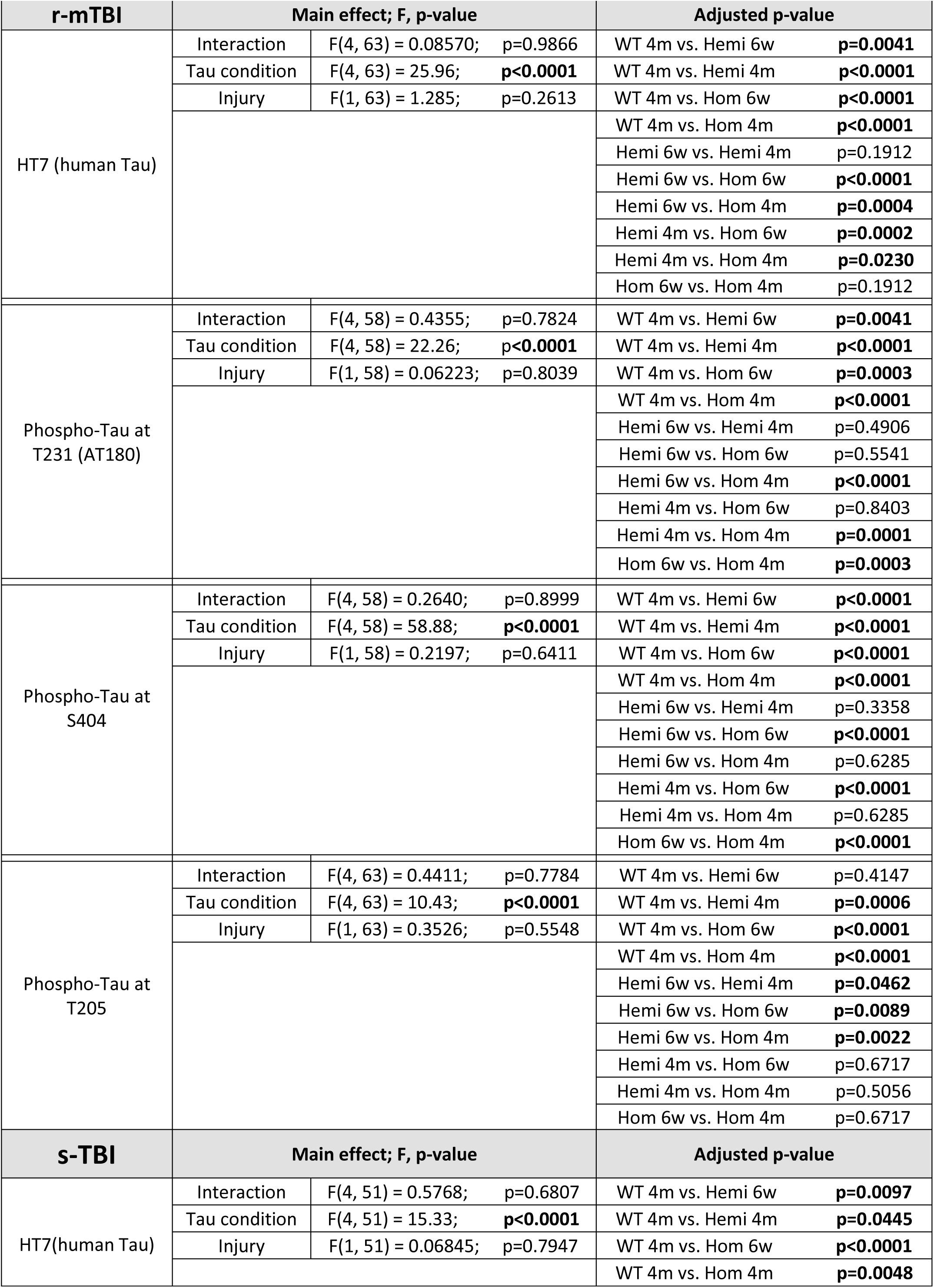
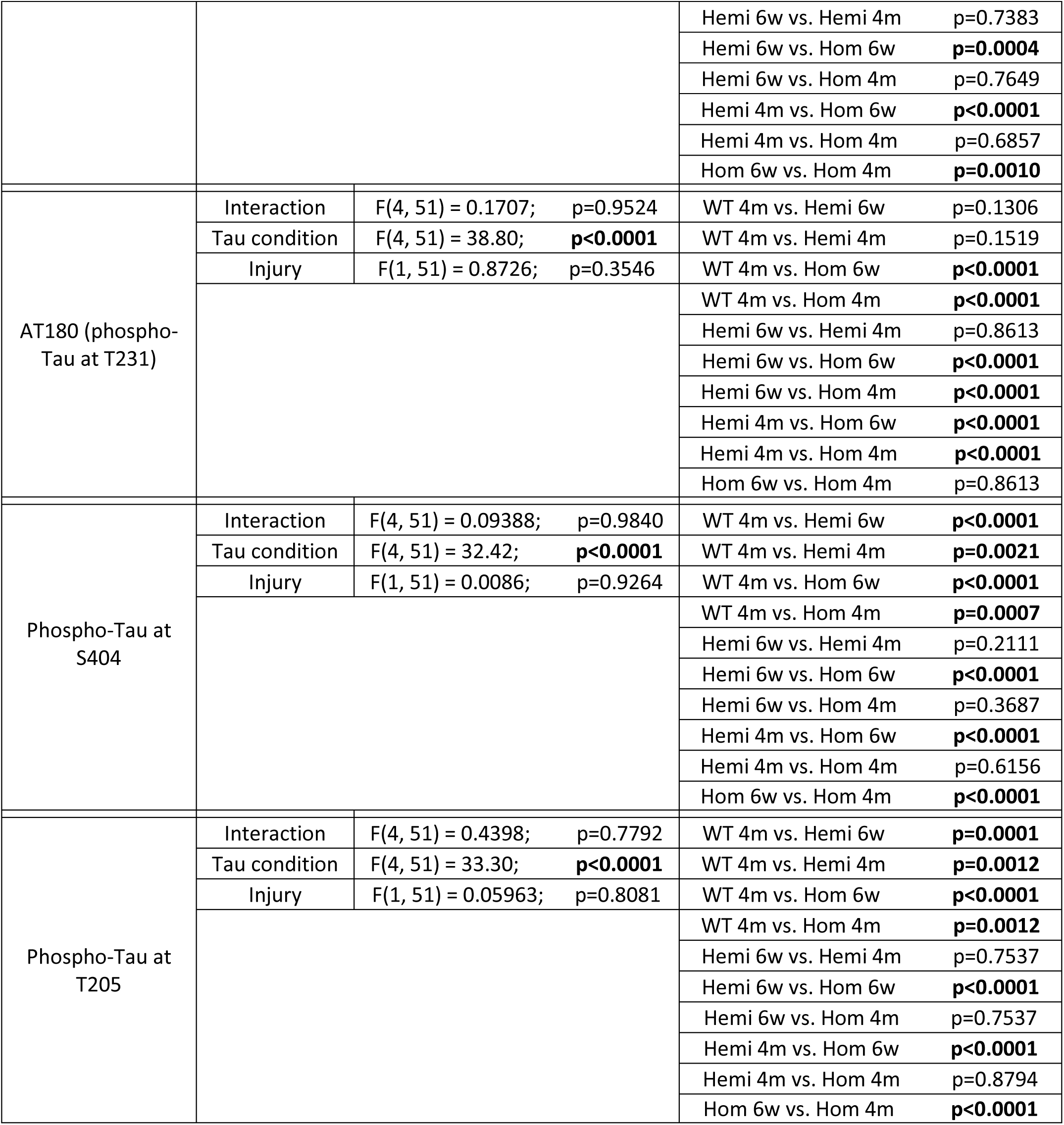
Wes for tau proteins in brain lysates from under impact site.

| **r-mTBI** | **Main effect; F, p-value** |  | **Adjusted p-value** |  |
| --- | --- | --- | --- | --- |
| HT7 (human Tau) | Interaction | F(4, 63) = 0.08570; p=0.9866 | WT 4m vs. Hemi 6w | **p=0.0041** |
|  | Tau condition | F(4, 63) = 25.96; **p<0.0001** | WT 4m vs. Hemi 4m | **p<0.0001** |
|  | Injury | F(1, 63) = 1.285; p=0.2613 | WT 4m vs. Hom 6w | **p<0.0001** |
|  |  |  | WT 4m vs. Hom 4m | **p<0.0001** |
|  |  |  | Hemi 6w vs. Hemi 4m | p=0.1912 |
|  |  |  | Hemi 6w vs. Hom 6w | **p<0.0001** |
|  |  |  | Hemi 6w vs. Hom 4m | **p=0.0004** |
|  |  |  | Hemi 4m vs. Hom 6w | **p=0.0002** |
|  |  |  | Hemi 4m vs. Hom 4m | **p=0.0230** |
|  |  |  | Hom 6w vs. Hom 4m | p=0.1912 |
| Phospho-Tau at T231 (AT180) | Interaction | F(4, 58) = 0.4355; p=0.7824 | WT 4m vs. Hemi 6w | **p=0.0041** |
|  | Tau condition | F(4, 58) = 22.26; **p<0.0001** | WT 4m vs. Hemi 4m | **p<0.0001** |
|  | Injury | F(1, 58) = 0.06223; p=0.8039 | WT 4m vs. Hom 6w | **p=0.0003** |
|  |  |  | WT 4m vs. Hom 4m | **p<0.0001** |
|  |  |  | Hemi 6w vs. Hemi 4m | p=0.4906 |
|  |  |  | Hemi 6w vs. Hom 6w | p=0.5541 |
|  |  |  | Hemi 6w vs. Hom 4m | **p<0.0001** |
|  |  |  | Hemi 4m vs. Hom 6w | p=0.8403 |
|  |  |  | Hemi 4m vs. Hom 4m | **p=0.0001** |
|  |  |  | Hom 6w vs. Hom 4m | **p=0.0003** |
| Phospho-Tau at S404 | Interaction | F(4, 58) = 0.2640; p=0.8999 | WT 4m vs. Hemi 6w | **p<0.0001** |
|  | Tau condition | F(4, 58) = 58.88; **p<0.0001** | WT 4m vs. Hemi 4m | **p<0.0001** |
|  | Injury | F(1, 58) = 0.2197; p=0.6411 | WT 4m vs. Hom 6w | **p<0.0001** |
|  |  |  | WT 4m vs. Hom 4m | **p<0.0001** |
|  |  |  | Hemi 6w vs. Hemi 4m | p=0.3358 |
|  |  |  | Hemi 6w vs. Hom 6w | **p<0.0001** |
|  |  |  | Hemi 6w vs. Hom 4m | p=0.6285 |
|  |  |  | Hemi 4m vs. Hom 6w | **p<0.0001** |
|  |  |  | Hemi 4m vs. Hom 4m | p=0.6285 |
|  |  |  | Hom 6w vs. Hom 4m | **p<0.0001** |
| Phospho-Tau at T205 | Interaction | F(4, 63) = 0.4411; p=0.7784 | WT 4m vs. Hemi 6w | p=0.4147 |
|  | Tau condition | F(4, 63) = 10.43; **p<0.0001** | WT 4m vs. Hemi 4m | **p=0.0006** |
|  | Injury | F(1, 63) = 0.3526; p=0.5548 | WT 4m vs. Hom 6w | **p<0.0001** |
|  |  |  | WT 4m vs. Hom 4m | **p<0.0001** |
|  |  |  | Hemi 6w vs. Hemi 4m | **p=0.0462** |
|  |  |  | Hemi 6w vs. Hom 6w | **p=0.0089** |
|  |  |  | Hemi 6w vs. Hom 4m | **p=0.0022** |
|  |  |  | Hemi 4m vs. Hom 6w | p=0.6717 |
|  |  |  | Hemi 4m vs. Hom 4m | p=0.5056 |
|  |  |  | Hom 6w vs. Hom 4m | p=0.6717 |
| **s-TBI** | **Main effect; F, p-value** |  | **Adjusted p-value** |  |
| HT7(human Tau) | Interaction | F(4, 51) = 0.5768; p=0.6807 | WT 4m vs. Hemi 6w | **p=0.0097** |
|  | Tau condition | F(4, 51) = 15.33; **p<0.0001** | WT 4m vs. Hemi 4m | **p=0.0445** |
|  | Injury | F(1, 51) = 0.06845; p=0.7947 | WT 4m vs. Hom 6w | **p<0.0001** |
|  |  |  | WT 4m vs. Hom 4m | **p=0.0048** |
|  |  |  | Hemi 6w vs. Hemi 4m | p=0.7383 |
|  |  |  | Hemi 6w vs. Hom 6w | **p=0.0004** |
|  |  |  | Hemi 6w vs. Hom 4m | p=0.7649 |
|  |  |  | Hemi 4m vs. Hom 6w | **p<0.0001** |
|  |  |  | Hemi 4m vs. Hom 4m | p=0.6857 |
|  |  |  | Hom 6w vs. Hom 4m | **p=0.0010** |
| AT180 (phospho-Tau at T231) | Interaction | F(4, 51) = 0.1707; p=0.9524 | WT 4m vs. Hemi 6w | p=0.1306 |
|  | Tau condition | F(4, 51) = 38.80; **p<0.0001** | WT 4m vs. Hemi 4m | p=0.1519 |
|  | Injury | F(1, 51) = 0.8726; p=0.3546 | WT 4m vs. Hom 6w | **p<0.0001** |
|  |  |  | WT 4m vs. Hom 4m | **p<0.0001** |
|  |  |  | Hemi 6w vs. Hemi 4m | p=0.8613 |
|  |  |  | Hemi 6w vs. Hom 6w | **p<0.0001** |
|  |  |  | Hemi 6w vs. Hom 4m | **p<0.0001** |
|  |  |  | Hemi 4m vs. Hom 6w | **p<0.0001** |
|  |  |  | Hemi 4m vs. Hom 4m | **p<0.0001** |
|  |  |  | Hom 6w vs. Hom 4m | p=0.8613 |
| Phospho-Tau at S404 | Interaction | F(4, 51) = 0.09388; p=0.9840 | WT 4m vs. Hemi 6w | **p<0.0001** |
|  | Tau condition | F(4, 51) = 32.42; **p<0.0001** | WT 4m vs. Hemi 4m | **p=0.0021** |
|  | Injury | F(1, 51) = 0.0086; p=0.9264 | WT 4m vs. Hom 6w | **p<0.0001** |
|  |  |  | WT 4m vs. Hom 4m | **p=0.0007** |
|  |  |  | Hemi 6w vs. Hemi 4m | p=0.2111 |
|  |  |  | Hemi 6w vs. Hom 6w | **p<0.0001** |
|  |  |  | Hemi 6w vs. Hom 4m | p=0.3687 |
|  |  |  | Hemi 4m vs. Hom 6w | **p<0.0001** |
|  |  |  | Hemi 4m vs. Hom 4m | p=0.6156 |
|  |  |  | Hom 6w vs. Hom 4m | **p<0.0001** |
| Phospho-Tau at T205 | Interaction | F(4, 51) = 0.4398; p=0.7792 | WT 4m vs. Hemi 6w | **p=0.0001** |
|  | Tau condition | F(4, 51) = 33.30; **p<0.0001** | WT 4m vs. Hemi 4m | **p=0.0012** |
|  | Injury | F(1, 51) = 0.05963; p=0.8081 | WT 4m vs. Hom 6w | **p<0.0001** |
|  |  |  | WT 4m vs. Hom 4m | **p=0.0012** |
|  |  |  | Hemi 6w vs. Hemi 4m | p=0.7537 |
|  |  |  | Hemi 6w vs. Hom 6w | **p<0.0001** |
|  |  |  | Hemi 6w vs. Hom 4m | p=0.7537 |
|  |  |  | Hemi 4m vs. Hom 6w | **p<0.0001** |
|  |  |  | Hemi 4m vs. Hom 4m | p=0.8794 |
|  |  |  | Hom 6w vs. Hom 4m | **p<0.0001** |

**Supplemental Information Table SI-6.**
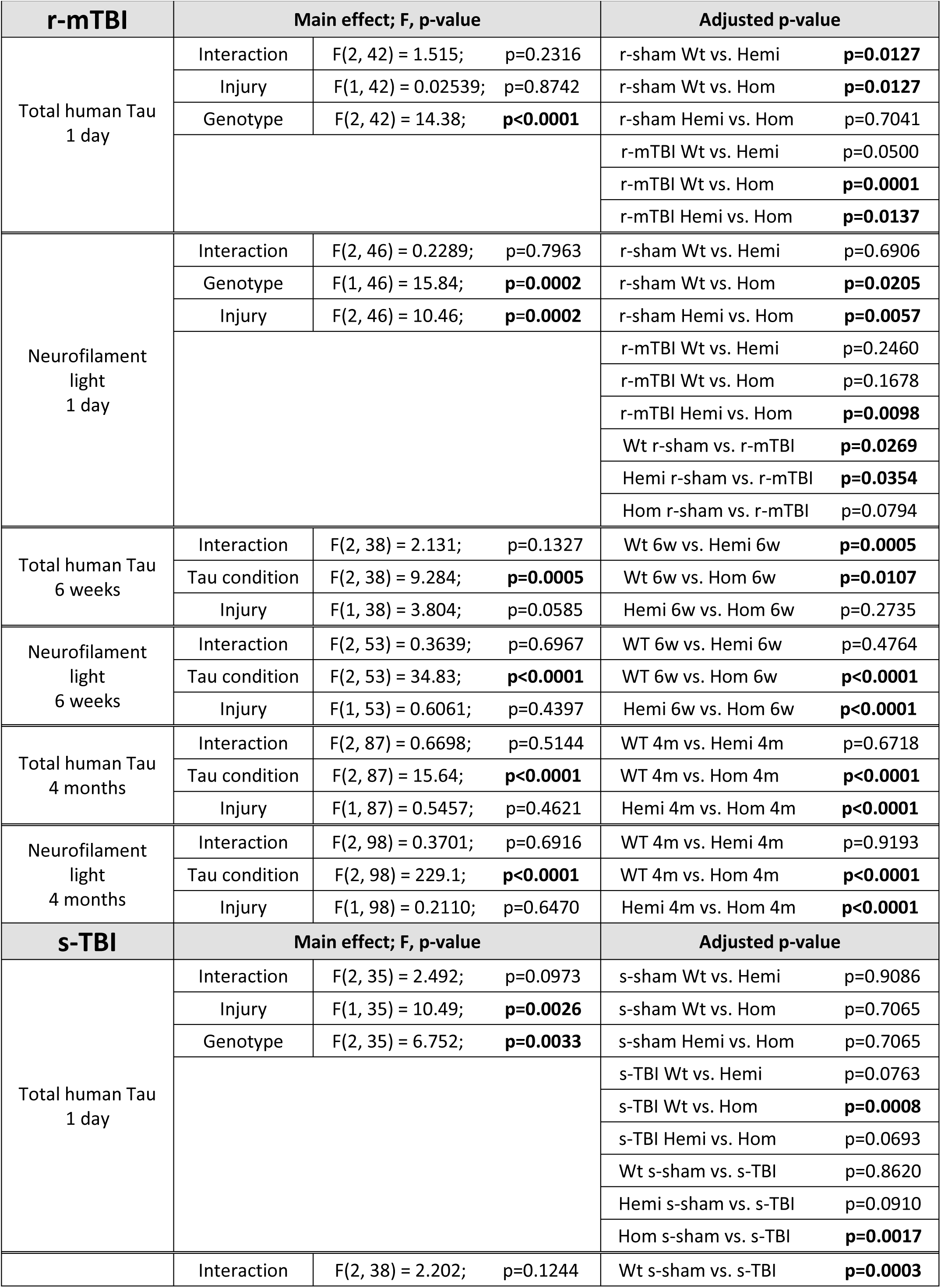
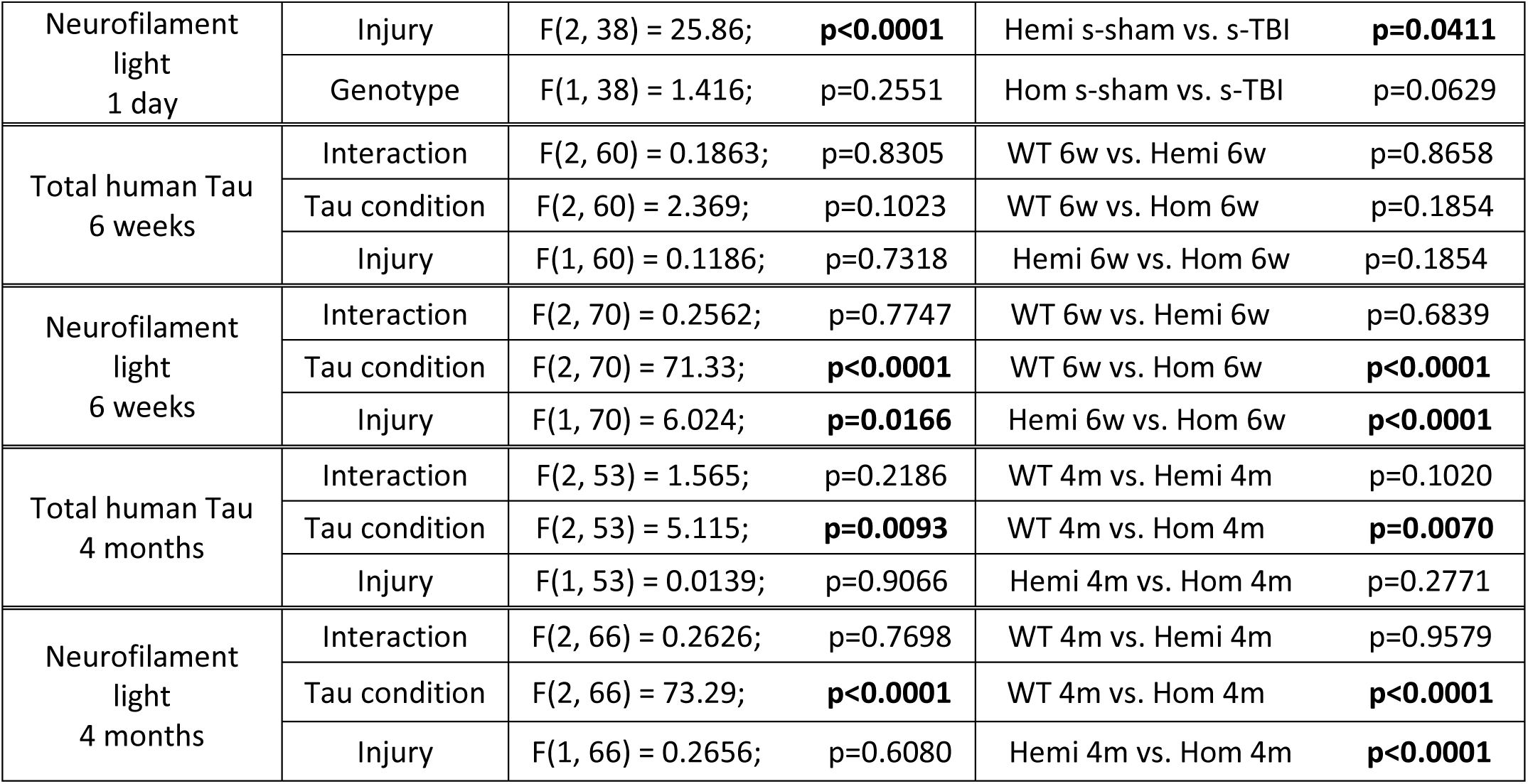
Simoa analysis of serum tau and neurofilament light protein.

| **r-mTBI** | **Main effect; F, p-value** |  |  | **Adjusted p-value** |  |
| --- | --- | --- | --- | --- | --- |
| Total human Tau<br>1 day | Interaction | F(2, 42) = 1.515; | p=0.2316 | r-sham Wt vs. Hemi | **p=0.0127** |
|  | Injury | F(1, 42) = 0.02539; | p=0.8742 | r-sham Wt vs. Hom | **p=0.0127** |
|  | Genotype | F(2, 42) = 14.38; | **p<0.0001** | r-sham Hemi vs. Hom | p=0.7041 |
|  |  |  |  | r-mTBI Wt vs. Hemi | p=0.0500 |
|  |  |  |  | r-mTBI Wt vs. Hom | **p=0.0001** |
|  |  |  |  | r-mTBI Hemi vs. Hom | **p=0.0137** |
| Neurofilament<br>light<br>1 day | Interaction | F(2, 46) = 0.2289; | p=0.7963 | r-sham Wt vs. Hemi | p=0.6906 |
|  | Genotype | F(1, 46) = 15.84; | **p=0.0002** | r-sham Wt vs. Hom | **p=0.0205** |
|  | Injury | F(2, 46) = 10.46; | **p=0.0002** | r-sham Hemi vs. Hom | **p=0.0057** |
|  |  |  |  | r-mTBI Wt vs. Hemi | p=0.2460 |
|  |  |  |  | r-mTBI Wt vs. Hom | p=0.1678 |
|  |  |  |  | r-mTBI Hemi vs. Hom | **p=0.0098** |
|  |  |  |  | Wt r-sham vs. r-mTBI | **p=0.0269** |
|  |  |  |  | Hemi r-sham vs. r-mTBI | **p=0.0354** |
|  |  |  |  | Hom r-sham vs. r-mTBI | p=0.0794 |
| Total human Tau<br>6 weeks | Interaction | F(2, 38) = 2.131; | p=0.1327 | Wt 6w vs. Hemi 6w | **p=0.0005** |
|  | Tau condition | F(2, 38) = 9.284; | **p=0.0005** | Wt 6w vs. Hom 6w | **p=0.0107** |
|  | Injury | F(1, 38) = 3.804; | p=0.0585 | Hemi 6w vs. Hom 6w | p=0.2735 |
| Neurofilament<br>light<br>6 weeks | Interaction | F(2, 53) = 0.3639; | p=0.6967 | WT 6w vs. Hemi 6w | p=0.4764 |
|  | Tau condition | F(2, 53) = 34.83; | **p<0.0001** | WT 6w vs. Hom 6w | **p<0.0001** |
|  | Injury | F(1, 53) = 0.6061; | p=0.4397 | Hemi 6w vs. Hom 6w | **p<0.0001** |
| Total human Tau<br>4 months | Interaction | F(2, 87) = 0.6698; | p=0.5144 | WT 4m vs. Hemi 4m | p=0.6718 |
|  | Tau condition | F(2, 87) = 15.64; | **p<0.0001** | WT 4m vs. Hom 4m | **p<0.0001** |
|  | Injury | F(1, 87) = 0.5457; | p=0.4621 | Hemi 4m vs. Hom 4m | **p<0.0001** |
| Neurofilament<br>light<br>4 months | Interaction | F(2, 98) = 0.3701; | p=0.6916 | WT 4m vs. Hemi 4m | p=0.9193 |
|  | Tau condition | F(2, 98) = 229.1; | **p<0.0001** | WT 4m vs. Hom 4m | **p<0.0001** |
|  | Injury | F(1, 98) = 0.2110; | p=0.6470 | Hemi 4m vs. Hom 4m | **p<0.0001** |
| **s-TBI** | **Main effect; F, p-value** |  |  | **Adjusted p-value** |  |
| Total human Tau<br>1 day | Interaction | F(2, 35) = 2.492; | p=0.0973 | s-sham Wt vs. Hemi | p=0.9086 |
|  | Injury | F(1, 35) = 10.49; | **p=0.0026** | s-sham Wt vs. Hom | p=0.7065 |
|  | Genotype | F(2, 35) = 6.752; | **p=0.0033** | s-sham Hemi vs. Hom | p=0.7065 |
|  |  |  |  | s-TBI Wt vs. Hemi | p=0.0763 |
|  |  |  |  | s-TBI Wt vs. Hom | **p=0.0008** |
|  |  |  |  | s-TBI Hemi vs. Hom | p=0.0693 |
|  |  |  |  | Wt s-sham vs. s-TBI | p=0.8620 |
|  |  |  |  | Hemi s-sham vs. s-TBI | p=0.0910 |
|  |  |  |  | Hom s-sham vs. s-TBI | **p=0.0017** |
|  | Interaction | F(2, 38) = 2.202; | p=0.1244 | Wt s-sham vs. s-TBI | **p=0.0003** |

|  |  |  |  |  |
| --- | --- | --- | --- | --- |
| Neurofilament<br>light<br>1 day | Injury | $F(2, 38) = 25.86$; **$p < 0.0001$** | Hemi s-sham vs. s-TBI | **$p = 0.0411$** |
| | Genotype | $F(1, 38) = 1.416$; $p = 0.2551$ | Hom s-sham vs. s-TBI | $p = 0.0629$ |
| Total human Tau<br>6 weeks | Interaction | $F(2, 60) = 0.1863$; $p = 0.8305$ | WT 6w vs. Hemi 6w | $p = 0.8658$ |
| | Tau condition | $F(2, 60) = 2.369$; $p = 0.1023$ | WT 6w vs. Hom 6w | $p = 0.1854$ |
| | Injury | $F(1, 60) = 0.1186$; $p = 0.7318$ | Hemi 6w vs. Hom 6w | $p = 0.1854$ |
| Neurofilament<br>light<br>6 weeks | Interaction | $F(2, 70) = 0.2562$; $p = 0.7747$ | WT 6w vs. Hemi 6w | $p = 0.6839$ |
| | Tau condition | $F(2, 70) = 71.33$; **$p < 0.0001$** | WT 6w vs. Hom 6w | **$p < 0.0001$** |
| | Injury | $F(1, 70) = 6.024$; **$p = 0.0166$** | Hemi 6w vs. Hom 6w | **$p < 0.0001$** |
| Total human Tau<br>4 months | Interaction | $F(2, 53) = 1.565$; $p = 0.2186$ | WT 4m vs. Hemi 4m | $p = 0.1020$ |
| | Tau condition | $F(2, 53) = 5.115$; **$p = 0.0093$** | WT 4m vs. Hom 4m | **$p = 0.0070$** |
| | Injury | $F(1, 53) = 0.0139$; $p = 0.9066$ | Hemi 4m vs. Hom 4m | $p = 0.2771$ |
| Neurofilament<br>light<br>4 months | Interaction | $F(2, 66) = 0.2626$; $p = 0.7698$ | WT 4m vs. Hemi 4m | $p = 0.9579$ |
| | Tau condition | $F(2, 66) = 73.29$; **$p < 0.0001$** | WT 4m vs. Hom 4m | **$p < 0.0001$** |
| | Injury | $F(1, 66) = 0.2656$; $p = 0.6080$ | Hemi 4m vs. Hom 4m | **$p < 0.0001$** |

## Notes

### Competing Interest Statement

The authors have declared no competing interest.

